# Comparing within- and between-family polygenic score prediction

**DOI:** 10.1101/605006

**Authors:** Saskia Selzam, Stuart J. Ritchie, Jean-Baptiste Pingault, Chandra A. Reynolds, Paul F. O’Reilly, Robert Plomin

**Author notes:** Corresponding Author Saskia Selzam.

## Abstract

Polygenic scores are a popular tool for prediction of complex traits. However, prediction estimates in samples of unrelated participants can include effects of population stratification, assortative mating and environmentally mediated parental genetic effects, a form of genotype-environment correlation (rGE). Comparing genome-wide polygenic score (GPS) predictions in unrelated individuals with predictions between siblings in a within-family design is a powerful approach to identify these different sources of prediction. Here, we compared within- to between-family GPS predictions of eight life outcomes (anthropometric, cognitive, personality and health) for eight corresponding GPSs. The outcomes were assessed in up to 2,366 dizygotic (DZ) twin pairs from the Twins Early Development Study from age 12 to age 21. To account for family clustering, we used mixed-effects modelling, simultaneously estimating within- and between-family effects for target- and cross-trait GPS prediction of the outcomes. There were three main findings: (1) DZ twin GPS differences predicted DZ differences in height, BMI, intelligence, educational achievement and ADHD symptoms; (2) target and cross-trait analyses indicated that GPS prediction estimates for cognitive traits (intelligence and educational achievement) were on average 60% greater between families than within families, but this was not the case for non-cognitive traits; and (3) this within- and between-family difference for cognitive traits disappeared after controlling for family socio-economic status (SES), suggesting that SES is a source of between-family prediction through rGE mechanisms. These results provide novel insights into the patterns by which rGE contributes to GPS prediction, while ruling out confounding due to population stratification and assortative mating.

## Introduction

The recent influx of well-powered genome-wide association (GWA) studies has led to substantial advances in our ability to detect genetic associations between single base pair variants (single nucleotide polymorphisms; SNPs) across the genome and a myriad of complex traits. Although individual SNP effect sizes are extremely small^1^, the surge in GWA power has improved the ability to predict complex traits through the genome-wide polygenic score (GPS) approach^2,3^. GPSs are indexes of individuals’ genetic propensity for a trait, and are derived as the sum of the total number of trait-associated alleles across the genome, weighted by their respective association effect size estimated through GWA analysis^4^. GPS can be calculated in any sample with genotype data that is independent from the discovery GWA study, and have permeated research in the social, behavioural and biomedical sciences^5^. In this paper, we use within-family analyses to investigate an important potential source of prediction in polygenic score analysis: passive genotype-environment correlation.

Currently one of the largest GWA meta-analyses with a sample size of 1.1 million was performed on educational attainment (years of schooling)^6^. A GPS derived from this study is the most predictive GPS for any behavioural trait to date, explaining 10.6% of the variance in years of education^6^ and 14.8% in tested educational achievement^7^. The predictive power of the educational attainment GPS (EA GPS) is considerable in contrast to other GPS for behavioral traits. Notably, cross-trait analyses have revealed that EA GPS is widely associated with traits other than educational achievement, including intelligence^2,6,7^, socioeconomic status (SES)^8–11^, behaviour problems^12^, mental illness^13^, physical health^13^ and personality^14,15^, in some cases accounting for as much as or more than the variance in cross-trait associations explained by the target GPS themselves^15,16^.

However, GWA analyses, and the GPSs derived from them in independent samples, are naïve to the pathways that lead from SNPs to trait outcomes^17^. With a focus on prediction, the mechanisms by which polygenic scores relate to phenotypes are left largely unexplored. Given the popularity and widespread use of the GPS approach, the interpretation of GPS prediction estimates requires more careful consideration. Potentially, *passive genotype-environment correlation* (prGE)^18^ effects could be one source of prediction. Parents generate family environments consistent with their own genotypes, which in turn facilitate the development of the offspring trait, thus inducing a correlation between offspring genotype and family environment^19–22^. Although these effects are also genetic in origin, they stem from the parents and are thus environmentally mediated. Therefore, GPS prediction among unrelated individuals may include contributions from both direct genetic effects and also indirect effects due to prGE.

Within-family analysis of siblings is a powerful approach to disentangle these potential sources of prediction. The additive genetic correlation between siblings is on average 0.50^23^, and the transmission of alleles from parents to offspring is randomized during meiosis, such that siblings have equal probability of inheriting any given allele^24^. The variability around the average genetic relationship between siblings due to random segregation is generally independent of the environment, therefore any genetic difference between siblings is free of shared environmental influence^25^. A relationship between their genetic differences and trait differences provides evidence for a causal effect of the measured genetic difference, since (i) siblings are well-matched on all shared familial genetic influences that shape the environment, and (ii) potential bias due to population stratification and assortative mating is completely eliminated within families^6,26,27^. Such within-family analyses account for prGE effects that are related to common family environments which are correlated with the transmitted alleles shared between siblings, but also environmental effects related to non-transmitted parental alleles that contribute to offspring similarity within a family. The use of DZ co-twins strengthens this design further as all shared environmental influences are time-invariant between twins (e.g. pregnancy risk factors, parental age, family income).

Indeed, previous within-family analyses have revealed substantial reductions in individual SNP effect sizes. For example, there was an effect size attenuation of ~40% compared to between-family associations in the most recent GWA study on educational attainment^6^. Most of this reduction has been attributed to prGE; no similar deflation of effect sizes was found for height^6^, indicating that prGE is not likely at play. A novel method relying on close- and distantly-related individuals, and that is applied to very large populations, detected a similar reduction of SNP-heritability estimates of educational achievement (35%)^25^. Moreover, studies that tested the effect between non-transmitted alleles from parental to offspring genotypes on offspring outcomes reported a significant association for educational attainment – an effect of so-called *genetic nurture* – but not for height and BMI^20,21^. In contrast, one study that tested within-family predictions of educational attainment using the EA GPS found no noteworthy difference in comparison to between-family estimates^28^. However, this GPS was based on the smallest GWA study for educational attainment^29^, and may have been underpowered to pick up prGE-driven effects.

Overall, relatively little research has been conducted on within-family GPS prediction and it has so far been limited to educational and anthropometric traits. This study adds substantially to this literature by systematically comparing within-family GPS prediction to between-family GPS prediction across eight core life outcomes (height, BMI, self-rated health, intelligence, educational achievement, neuroticism, attention-deficit/hyperactivity symptoms, and schizophrenia symptoms). Educational achievement is both phenotypically and genetically correlated with many life outcomes^30–36^. It is also is highly genetically correlated with family SES^8,37,38^, and EA GPS predicts 7.3% of the variance in SES^9^. Therefore, it is possible that the effects identified in the GWA studies for educational attainment related to family environment (e.g. SES) also contribute to the development of other behavioural traits through prGE mechanisms. Although it has been suggested that the widespread cross-trait associations between the EA GPS and various outcomes may be partly driven by prGE effects^15,22^, to our knowledge no study to date has tested this hypothesis.

It is the aim of this study to investigate potential influences of prGE in a range of life outcomes through the comparison of within- and between-family polygenic score prediction estimates. First, we predict that within-family estimates will be disproportionally lower than between-family estimates for EA GPS predictions of educational achievement in contrast to other GPS predictions of their target trait. Second, we predict that cross-trait associations between the EA GPS and other outcomes will be smaller within families than between families, in comparison to the cross-trait associations of other GPS.

## Methods

Our hypotheses, measures and analysis plan were preregistered with the Open Science Framework (for more details, see Online Resource section), except where indicated below. The non-preregistered analyses should be considered exploratory.

### Sample

Participants were drawn from the Twins Early Development Study (TEDS). Between 1994-1996 TEDS recruited 16,810 twin pairs born in England and Wales, who have been assessed in multiple waves across development until the present. The demographic characteristics of TEDS participants and their families closely match those of families in the UK^9,39^. Written informed consent was obtained from parents prior to data collection, and from TEDS participants themselves past the age of 18. Project approval was granted by King’s College London’s ethics committee for the Institute of Psychiatry, Psychology and Neuroscience PNM/09/10–104. Only DZ co-twins with complete data were included in this study.

### Phenotypic data

#### Height

Self-reported height was assessed at the average age of 22.1 (SD=0.86) in 1,463 twin pairs, including 789 same-sex and 674 opposite-sex twin pairs.

#### Body Mass Index (BMI)

BMI was calculated using self-reported weight in kg and height in meters 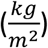 at age 22.1 (SD=0.86) in 1,353 twin pairs, including 733 same-sex and 620 opposite-sex twin pairs.

#### Self-rated health

Twins rated their health on the reduced RAND Short-Form Health Survey^40^. Individuals scored their health on a five point Likert scale for questions such as “In general, would you say your health is?” (“Poor” to “Excellent”), or “I am as healthy as anybody I know” (“Strongly Disagree” to “Strongly Agree”). Data were available on 1,494 twin pairs, including 805 same-sex and 689 opposite-sex twin pairs at age 22.1 (SD=0.86).

#### Intelligence

At age 11.4 (SD=0.65), twins were assessed on their non-verbal abilities (Raven’s Standard Progressive Matrices^41^; WISC-III-UK Picture Completion^42^) and on their verbal abilities (WISC-III-PI Vocabulary Multiple-Choice^43^; WISC-III-PI Information Multiple-Choice^43^). A composite variable was calculated as the arithmetic mean of the z-standardized scales for 1,569 twin pairs, including 824 same-sex and 745 opposite-sex twin pairs.

#### Educational achievement

Results for standardized tests taken at the end of compulsory education in the United Kingdom (General Certificate of Secondary Education; GCSE) were obtained for twins at age 16.3 (SD=0.29) via self-report. Grades were coded from 4 (G; the minimum pass grade) to 11 (A* the highest possible grade). Self-reported GCSE grades in TEDS highly correlate with grades obtained for a subsample of individuals from the National Pupil Database (*r* = 0.98 for English, *r* = 0.99 for mathematics, *r* > 0.95 for all sciences)^31^. A composite was calculated as the arithmetic mean of the compulsory core subjects – Maths, English and Science – for 2,366 twin pairs, including 1,220 same-sex and 1,146 opposite-sex twin pairs.

#### Neuroticism

At age 16.5 (SD=0.27), twins were assessed on their Big Five personality traits on a five-point Likert scale^44^. For this study, we used the six Neuroticism items (e.g. Anxiousness; Vulnerability) to form a composite score by taking the arithmetic mean for 789 twin pairs, including 429 same-sex and 360 opposite-sex twin pairs.

#### Attention-Deficit Hyperactivity Disorder (ADHD) symptoms

At age 11.5 (SD=0.69) and 16.3 (SD=0.69), parents reported on twins’ ADHD symptoms via the Strength and Difficulties Questionnaire^45^ hyperactivity subscale (three-point Likert scale) and the Conners’ rating scales (CPRS-R; four-point Likert scale)^46^ on hyperactivity and inattention. Although self-report ratings were available, it has been shown that informant-based ratings are more reflective of objective measures of ADHD symptoms^47^. A composite score was created as the arithmetic mean of the sex and age z-standardized scales. Where ratings were available at one assessment only, this value was used to maximise sample size, leading to a sample size of 2,469 twin pairs, including 1,285 same-sex and 1,184 opposite-sex twin pairs.

#### Schizophrenia symptoms

At age 22.7 (SD=0.85), paranoia and hallucinations were assessed through self-reported ratings on the Specific Psychotic Experiences Questionnaire (SPEQ; six point Likert scale)^47^, and parent-reported negative symptoms using the Scale for the Assessment of Negative Symptoms (SANS; four point Likert scale)^7^. Data were available for 1,140 twin pairs, including 613 same-sex and 527 opposite-sex twin pairs.

#### Family socio-economic status (SES)

This measure was calculated as the mean of the z-standardized maternal age at birth of the first child, maternal and paternal highest education level (coded from 1 = “no qualifications” to 8 = “postgraduate qualifications”), and maternal and paternal occupation (coded from 1 = “Other Occupations – dockers, porters, labourers,…” to 9 = “Managers and Administrators”). These measures were assessed at first contact at age 1.8 (SD=1.13). Data were available for 2,962 twin pairs, including 1,542 same-sex and 1,420 opposite-sex twin pairs.

Measures were selected based on largest sample sizes available, and ages at phenotype assessment matching most closely the ages of GWA study samples to maximise predictive power. None of the measures were significantly associated with birth order, but most showed sex and age differences (see Supplementary Table S1) and were therefore adjusted for these effects using the regression method, and z-standardised residuals (mean=0, SD=1) were used for all subsequent analyses.

### Genotypic data

Two different genotyping platforms were used because genotyping was undertaken in two separate waves, five years apart. AffymetrixGeneChip 6.0 SNP arrays were used to genotype 3,665 individuals. Additionally, 8,122 individuals (including 3,607 dizygotic co-twin samples) were genotyped on HumanOmniExpressExome-8v1.2 arrays. After quality control, 635,269 SNPs remained for AffymetrixGeneChip 6.0 genotypes, and 559,772 SNPs for HumanOmniExpressExome genotypes.

Genotypes from the two platforms were separately phased and imputed into the Haplotype Reference Consortium (release 1.1) through the Sanger Imputation Service^6^ before merging. Genotypes from a total of 10,346 samples (including 3,320 dizygotic twin pairs and 7,026 unrelated individuals) passed quality control, including 3,057 individuals genotyped on Affymetrix and 7,289 individuals genotyped on Illumina. The final data contained 7,363,646 genotyped or well imputed SNPs (for full genotype processing and quality control details, see^6^). To ease high computational demands of the software that generates polygenic scores, we further excluded SNPs with info <1, leaving 515,000 SNPs for analysis.

We performed principal component analysis on a subset of 39,353 common (MAF > 5%), perfectly imputed (info = 1) autosomal SNPs, after stringent pruning to remove markers in linkage disequilibrium (r2 > 0.1) and excluding high linkage disequilibrium genomic regions so as to ensure that only genome-wide effects were detected.

### Polygenic scores

We calculated polygenic scores based on summary statistics for the largest GWA studies available for key developmental outcomes, including height^48^, body mass index (BMI)^48^, self-rated health^45^, intelligence^47^, educational attainment^6^, neuroticism^49^, ADHD^50^, and schizophrenia^51^. These GWA studies were selected because their respective GPS yield the highest predictive accuracy within their trait category (details about the studies, reported SNP heritabilities and GPS predictions can be found in Supplementary Table S2). All polygenic scores were statistically adjusted for the first ten principal components, chip and plate using the regression method and were z-standardized (mean=0, SD=1).

### Polygenic scoring method

To calculate polygenic scores, we used a Bayesian approach, *LDpred*^52^. Using LDpred, a posterior effect size for each SNP is derived by re-weighting the original summary statistic coefficient based on (i) the relative influence of a SNP given its level of LD with surrounding SNPs, and (ii) a prior on the effect size of each SNP. This prior is dependent on the heritability of the trait, as well as the fraction of markers assumed to causally influence the trait. The final GPS is obtained as the sum of the trait-increasing alleles (each variant coded as 0, 1, or 2), weighted by the posterior effect size estimates. In contrast to clumping and thresholding, LDpred retains all the SNPs in the polygenic score that are common between GWA summary statistics and genotype data in the target sample (for more details about this polygenic score calculation approach, see^15^). In this study, we created polygenic scores using a prior that assumes a causal fraction of 1 for all analyses, based on the assumption that all genetic markers contribute to trait development.

## Statistical Analysis

### Mixed-effects modelling

We applied a mixed-effects model on DZ data with the novel approach of including two fixed effects to separate the total effect between the polygenic score predictor and the outcome into within- and between-family effects^53^:

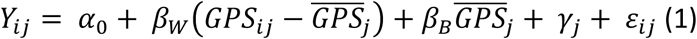

where *Y* denotes the outcome and GPS the polygenic score, *i* = {1,2} corresponds to the individual twins that are clustered within family *j*, and 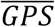 refers to the mean GPS value in family *j*. The *i*th value represents birth order, where twin 1 is the elder twin. The notation *α*_0_ represents the intercept and *γ*_*j*_ the random effect with 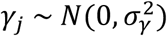, which corresponds to a change in the intercept for both twins in family *j*, and *ε*_*ij*_ with 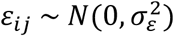, which denotes the independent random error for each individual *i* in family *j*. The between-family effect *β*_*B*_ represents the expected change in the outcome *Y* given a one unit change in the family GPS average, and the within-family effect *β*_*W*_ represents the expected change given a one unit change in the difference between the individual GPS and the family average GPS. By including both *β*_*W*_ and *β*_*B*_ in the same model, the individual estimates are adjusted for, and independent of, the effect of the other estimate. The random effect term 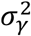, which estimates the difference between each group intercept *γ*_*j*_ and the overall intercept *α*_0_, accounts for the residual structure in the data corresponding to all unaccounted familial factors (both genetic and environmental) that contribute to the trait similarity of the twins^54,55^.

The use of a mixed-effects model is only justified if co-twins within a family correlate in the outcome, which can be estimated through the Intraclass Correlation (ICC) coefficient. The ICC is the ratio of the between-family (i.e. random intercept) variance over the total variance and is an estimate of how much of the total variation in the outcome is accounted for by family:

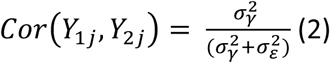

where 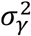 is the covariance between the family variable, in this case family ID, and the outcome, and 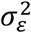 indicates the residual variance capturing within twin pair differences. The total effect of the relationship between GPS and outcome is the ICC weighted sum of the within- and between-family effects^53^:

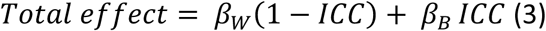

It follows from (3) that the total effect ranges between *β*_*W*_ and *β*_*B*_. If the relationship between GPS and outcome is mostly due to individual-level variation, the ICC approximates 0 and the total effect will be close to *β*_*W*_. In contrast, if the association is mostly due to family effects, the ICC approximates 1 and the total effect will be close to *β*_*B*_ ^47^.

Performing a regression corresponding to equation (1), we estimated the *β*_*W*_ and *β*_*B*_ parameters using each of the eight polygenic scores in turn as predictors of each of the eight measured outcomes. To estimate potential SES effects, we repeated these analyses including the SES composite as a covariate in the model (these latter analyses were not preregistered).

To empirically test the statistical difference between *β*_*W*_ and *β*_*B*_, we performed permutation testing. We randomly permuted the vector containing GPS values of twin 2, so that there was no longer a systematic relationship between the GPS of twin 1 and twin 2, in line with the null hypothesis of interchangeability. Model (1) was repeated 100,000 times using randomly sampled data with replacement, calculating the difference between *β*_*B*_ and *β*_*W*_ for each iteration, therefore creating an empirical null distribution. To test the null hypothesis, an empirical *p*-value was calculated as the fraction of the total number of permuted (i.e. random) difference values that were at least as extreme as the difference between *β*_*B*_ and *β*_*W*_ derived from unpermuted data. A pseudo-count of 1 was added to both numerator and denominator to avoid *p*-values of zero^56^, which would occur if all permuted difference values were smaller than the original difference in betas. Therefore, the lowest *achievable p*-value was 9.99e^−6^. Differences between *β*_*B*_ and *β*_*W*_ were considered as statistically significant if the empirical *p*-value was smaller than the alpha significance threshold (see below for multiple testing correction), indicating that this difference was unlikely a result of random sample characteristics.

### Quantile analysis of within-DZ pair differences

To illustrate the extent to which within-DZ pair GPS differences result in differences in developmental outcomes, we performed quantile analysis. Firstly, we generated twin-GPS difference scores by subtracting the twin 2 score from the twin 1 score, and then split this variable into ten equal quantiles based on absolute GPS differences, ranging from the lowest to the highest GPS differences. Birth order did not explain any statistically significant amount of variance (Supplementary Table S1), therefore no randomisation of twin order was required. We tested mean differences in outcome variables between individuals in the lowest and highest decile. We performed quantile analysis on variables with scales that are easily interpretable: that is, BMI, height, intelligence and educational achievement. For this purpose, the z-standardised and cleaned variables were transformed back to their original scale, and intelligence values were scaled to have a mean of 100 and a standard deviation of 15.

### Multiple testing correction

Multiple testing correction of the α significance threshold was performed using the Benjamini Hochberg false discovery rate (FDR) adjustment^57^. In contrast to more conservative corrections, this method has higher statistical power to detect true positives while controlling for false positives. Based on an α threshold of 0.05, the corrected α in this study was 0.01, defined as the maximum raw *p*-value that is smaller or equal than the FDR critical value 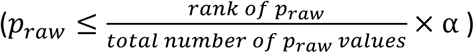.

## Results

Phenotypic resemblance between DZ twins within a family varied between traits, with Pearson’s correlation coefficients ranging from 0.10 – 0.59 (Supplementary Figure S1, and Supplementary Table S3 for ICCs). Twins were least alike in their neuroticism levels and self-rated health, and most alike in their height, IQ and educational achievement. Within twin pair polygenic score correlations were close to expectations (range *r* = 0.49 – 0.57), given that the expected shared additive genetic variance between siblings is 50% of the total additive genetic variance based on quantitative genetic theory^23^.

### Within-family polygenic score predictions

Figure 1 depicts the within- and between-family polygenic score prediction estimates of the eight core developmental outcomes from the mixed-effects model analyses. Within-family target-trait predictions were statistically significant for height, BMI, intelligence, educational achievement and ADHD symptoms, indicating that polygenic variation within twin pairs was related to these outcome differences. Specifically, phenotypic differences in height were significantly positively correlated with height GPS twin differences (*β* = 0.41, *p* = 5.72e^−53^) and differences in BMI were significantly correlated with BMI GPS differences (*β* = 0.30, *p* = 1.76e^−21^) such that twins with a higher height GPS and BMI GPS were taller and heavier than their co-twin, respectively. IQ GPS differences predicted intelligence differences (*β* = 0.14, *p* = 1.32e^−6^) and EA GPS differences were significantly associated with GCSE grade differences (*β* = 0.21, *p* = 2.22e^−26^), indicating that those twins with a higher GPS also scored higher on intelligence measures and in their GCSE tests than their co-twin. For behaviour problems, twins with higher ADHD GPS had higher phenotypic ADHD symptoms than their co-twins (*β* = 0.12, *p* = 1.50e^−7^).

**Figure 1.**
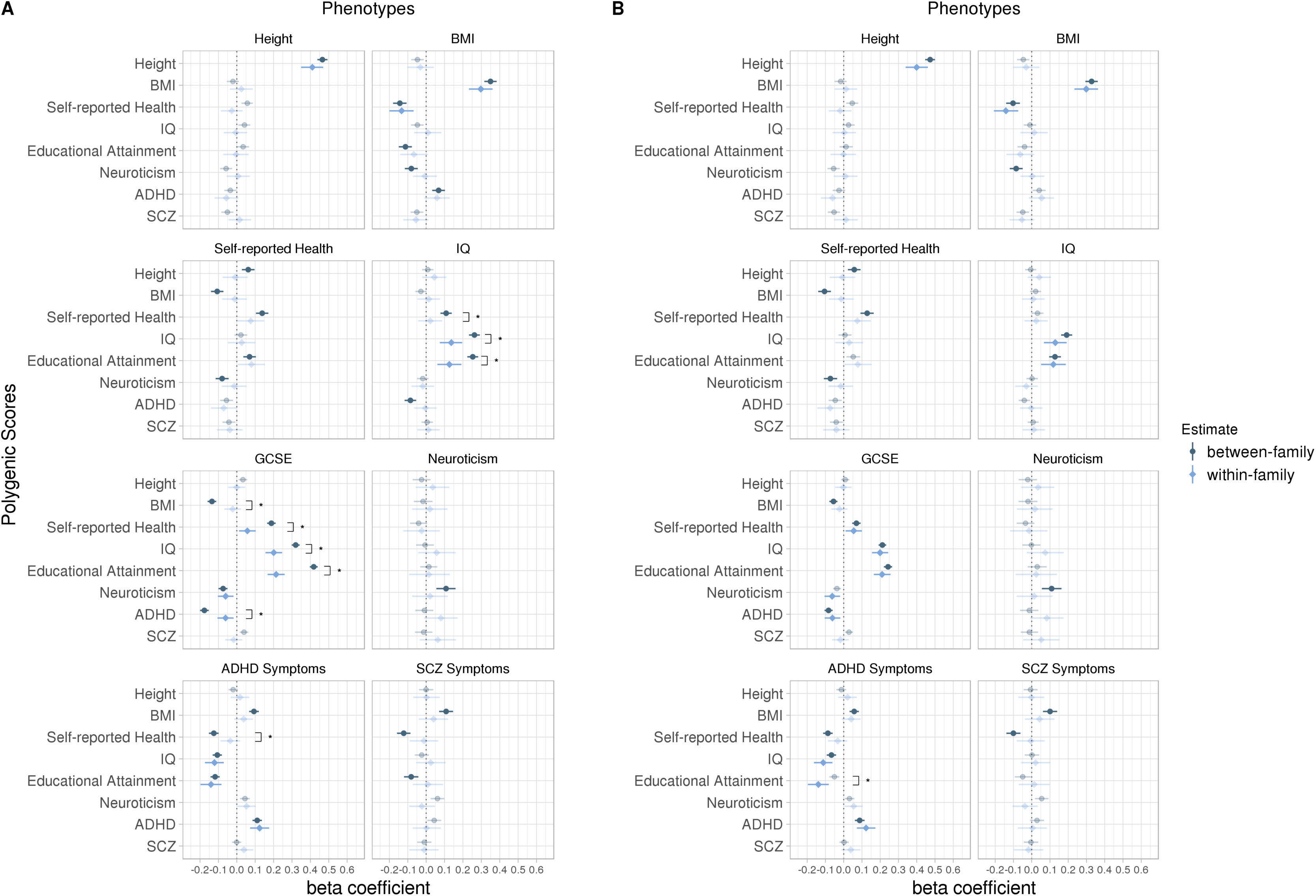
Within- and between-family prediction estimates of eight developmental outcomes using eight genome-wide polygenic scores (GPS) before (A) and after (B) statistical correction for family socio-economic status. The GPS are presented on the y-axis, predicting each of the eight phenotypic traits. Error bars are 95% bootstrap percentile intervals based on 10,000 bootstrap samples. Opaque estimates indicate statistical significance at the false discovery rate corrected threshold of *p* < 0.01. Brackets indicate a significant difference between within- and between-family prediction estimate based on permutation testing with 100,000 iterations. significant differences are only shown where at least one of the estimates is statistically significant at the false discovery rate corrected threshold of *p* < 0.01 (for all prediction estimates and *p*-values, see Supplementary Tables S4 and S5). The dotted line represents a beta coefficient of zero. BMI = Body Mass Index; IQ = Intelligence; GCSE = General Certificate of Secondary Education (educational achievement); ADHD = Attention-Deficit/Hyperactivity Disorder; SCZ = Schizophrenia.

We also investigated cross-trait relationships (Figure 1). For example, self-rated health GPS differences were negatively correlated with differences in BMI, such that twins with a higher self-rated health GPS had a lower BMI (β = − 0.13, p = 3.56e^−5^). EA GPS differences significantly related to phenotypic intelligence differences (β = 0.13, p = 2.15e^−5^), and IQ GPS predicted GCSE grade differences (β = 0.20, p = 7.24e^−25^), suggesting that those with higher GPS also had higher IQ and GCSE grades than their co-twin. GCSE grade differences were also negatively predicted by ADHD GPS twin differences (*β* = − 0.07, *p* = 2.20e^−4^), indicating that twins with a higher ADHD GPS obtain lower GCSE results. Notably, IQ GPS differences (*β* = − 0.12, *p* = 6.38e^−7^) and EA GPS differences (*β* = − 0.14, *p* = 3.09e^−8^) were just as predictive of ADHD symptoms as the ADHD GPS itself, and the direction of effect sizes indicates that the twin with a higher GPS had lower ADHD symptoms than their co-twin (all prediction estimates and total effects are presented in Supplementary Table S4).

### Comparing within-family and between-family polygenic score prediction

By simultaneously and independently estimating within- and between-family GPS predictions, it was possible to compare these estimates. Between-family estimates (Figure 1) are mostly consistent with GPS correlations reported for unrelated individuals (Supplementary Table S2). Figure 1 also shows that between-family associations are generally greater than within-family associations. Significant associations were found for 46.9% of the between-family associations and only 20.3% for within-family associations. On average, magnitudes of within-family associations were almost half (44.1% reduction) that compared to significant between-family estimates (for all prediction estimates and significance of differences see Supplementary Table S4).

Notably, significant differences in associations within and between families for polygenic scores predicting their target traits were almost exclusively found for IQ and educational achievement (Figure 1A). The within-family prediction was significantly lower than between-family prediction for both IQ (*p* = 3.50e^−4^, Δ = 48.0%) and GCSE grades (*p* = 9.99e^−6^, Δ = 48.9%).

Also, for cross-trait associations, differences in within- and between-family polygenic score predictions were most pronounced for IQ and educational achievement. For IQ, there were significant differences for the self-rated health GPS (*p* = 8.00e^−3^, Δ = 79.4%) and the EA GPS (*p* = 6.00e^−5^, Δ = 50.1%). For educational achievement, there were significant differences for the BMI GPS (*p* = 9.99e^−6^, Δ = 83.3%), the self-rated health GPS (*p* = 9.99e^−6^, Δ = 69.5%), the IQ GPS (*p* = 9.99e^−6^, Δ = 37.2%), and the ADHD GPS (*p* = 2.00e^−5^, Δ = 65.4%). In addition, there was a significant difference in within- and between-family prediction for the self-rated health GPS (*p* = 3.90e^−4^, Δ = 71.7%) predicting ADHD symptoms.

The finding that polygenic score prediction estimates of our measured traits are substantially smaller within families suggests that the corresponding between-family associations are mediated by some combination of family-specific (i.e. shared family) effects, population stratification and potentially assortative mating. Family SES, which is the same for members of a family, is a predictor not only of educational achievement and IQ, but also physical and mental health outcomes. Therefore, we repeated our analyses including family SES as a covariate in the model to interrogate its role in between-family GPS prediction. As noted above, this analysis was not pre-registered. As shown in Figure 1B, between-family predictions were greatly reduced and magnitudes approached those of within-family prediction estimates, which did not change (because any shared family effects are already controlled for in within-family estimates; for all prediction estimates and significance of differences, see Supplementary Table S5). For IQ, educational achievement, and ADHD, where the greatest prediction differences were observed, the average of statistically significant between-family beta estimates versus within-family estimates was 0.182 and 0.097, and 0.102 and 0.095 after accounting for SES, respectively.

Sensitivity analyses (not pre-registered) were performed by repeating all analyses separately for same-sex and opposite-sex twins, and we found no substantial deviations from the results using the combined sample (For results, see Supplementary Tables S6 to S10, and Supplementary Figures S2 and S3).

### Quantile Analysis

To illustrate within-family differences further, quantile analysis demonstrated how within-family polygenic score differences related to differences in height, BMI, intelligence and GCSE grades (Figure 2). There was an 8.7cm height mean difference (*p* = 1.28e^−11^) between the lowest absolute difference decile versus the highest difference decile. For BMI, the difference was 2.9 BMI points (*p* = 8.33e^−6^) between the lowest and the highest absolute GPS difference deciles. Mean GCSE grade differences (0.40) were also statistically significant (*p* = 7.13e^−5^) when comparing the lowest and the highest absolute GPS difference deciles. In contrast, intelligence point differences (1.9 points) were not statistically different (*p* = 0.26) between the lowest and the highest absolute GPS difference quantiles (for trait and GPS means at each difference decile see Supplementary Table S11).

**Figure 2.**
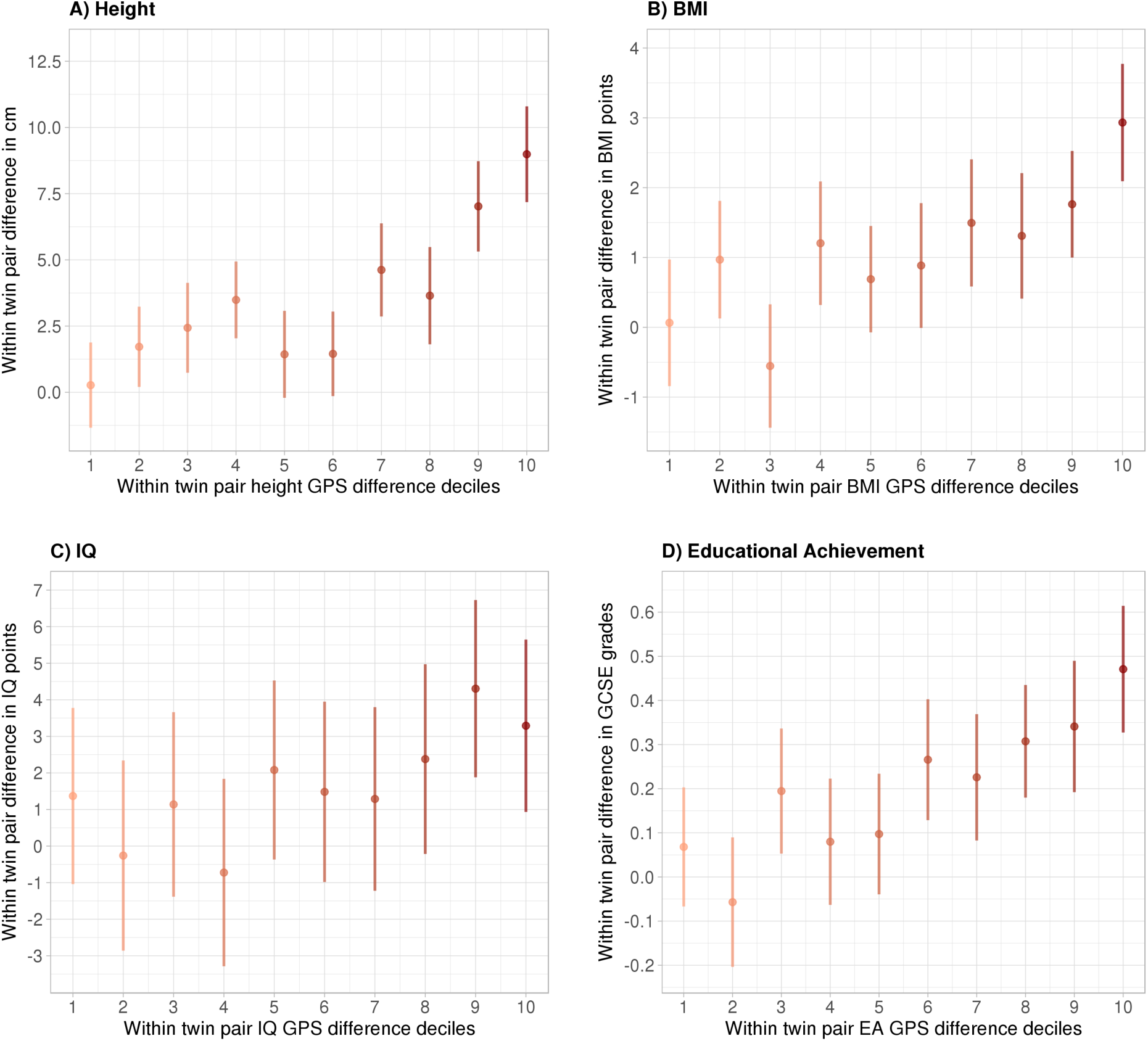
The relationship between absolute DZ twin polygenic score decile differences and trait outcome differences. Lower deciles represent small absolute genome-wide polygenic score (GPS) differences and higher deciles represent large GPS differences between DZ co-twins. Error bars indicate 95% confidence intervals. Each GPS decile included the following numbers of twin pairs: Height = 146; BMI = 135; IQ = 157; GCSE = 236. Regression through origin analysis (fixed intercept of zero) using the continuous GPS difference values to predict outcome differences were significant for height (*B*= 4.42, *p* = 3.73e^−53^, *R*^2^ = 0.148), BMI (*B*= 1.34, *p* = 1.73e^−21^, *R*^2^ = 0.064), IQ (*B*= 2.1, *p* = 4.53e^−7^, *R*^2^ = 0.015), and GCSE grades (*B*= 0.26, *p* = 3.04e^−26^, *R*^2^ = 0.046).

## Discussion

Polygenic score prediction of complex traits is now a common approach in genomics research, but the potential pathways by which polygenic score variation predicts phenotypic variation remain largely unexplored. In this study, we contrasted within- and between-family polygenic prediction estimates to quantify the extent to which environmentally-mediated genetic effects (i.e. passive genotype-environment correlation) are picked up in polygenic score analyses. By systematically performing target- and cross-trait analyses across eight life outcomes using eight corresponding GPS, we found evidence that prGE might be a mechanism explaining a considerable proportion of the GPS prediction in cognitive traits (intelligence and educational achievement), but not for non-cognitive traits. We also found that for all between-family GPS predictions of cognitive traits – but, again, not other traits – family SES is likely to be the major source of prGE.

For the prediction of IQ and educational achievement, within-family estimates were on average 60% smaller than between-family estimates. The within-versus between-family attenuation for the EA GPS prediction was 49%, which is close to the 40% estimate in GWA study effect sizes for years of education^6^. These findings highlight the influence of prGE in the development of IQ and educational achievement, and demonstrate the extent to which between-family GPS prediction may be partly driven by prGE effects. Results from our study are also in line with adoption studies showing evidence of between-family prGE in that correlations between home environment and children’s IQ is twice as great in non-adoptive families than in adoptive families^58^. Our findings are compatible with recent research on *genetic nurture*, using non-transmitted alleles from parental genotypes to assess prGE^20,21^ in terms of GPS target trait prediction of educational achievement and anthropometric traits. Our findings are novel to the extent that we performed cross-trait associations using a wide range of GPS. Contrary to our prediction that within- and between-family EA GPS associations would be significantly different across many associated life outcomes, results from cross-trait analysis suggest that within- and between-family predictions were only significantly different across a range of GPS for the prediction of cognitive outcomes.

A possible explanation for these results is that IQ and educational achievement show more shared environmental influences (24% and 27%, respectively) relative to other traits used in this study such as height (10%), BMI (10%), ADHD (2%), or schizophrenia (0%), as estimated through a large twin study meta-analysis^59^. The type of rGE that we assessed in this study – defined as the exposure to a family environment that is correlated with both parental and offspring genotypes, and which contributes to sibling similarity in their outcomes – is absorbed by the shared environment variance component (‘C’) in classical twin analyses^60^. Therefore, it may be more likely that genetic effects related to cognitive traits as estimated through GWA studies partly contain prGE effects – in contrast to other traits tested in our study – because the shared environmental component is larger to begin with for cognitive traits. As known from the existing literature, family SES is strongly genetically correlated with offspring cognitive traits^8,37,38^, rendering it a likely source of prGE. Indeed, our results showed that between-family effects were similar in magnitude to within-family effects in magnitude when holding SES constant, suggesting that SES is a source of the majority of the within-between discrepancy, rather than residual population stratification or assortative mating.

The results showed that more distantly-related GPS captured considerable prGE effects in cross-trait GPS predictions of cognitive traits. For instance, within-family effect sizes for the ADHD GPS predicting educational achievement were significantly smaller (65% reduction), in contrast to the ADHD GPS predicting ADHD symptoms, where no significant difference was detected. This suggests that the GWAS for ADHD captures genetic variation that is correlated with aspects of the family environment that contribute to the co-development of ADHD symptoms and educational achievement, although it is unclear why these effects do not appear to contribute to the development of ADHD symptoms themselves.

It is important to go beyond GPS predictions of traits in unrelated individuals to consider prGE mechanisms by comparing within- and between-family predictions in order to explain the sources of predictions in polygenic score analysis. However, finding between-family prGE does not diminish the usefulness of GPS predictions for cognitive traits in unrelated individuals, because these prGE effects help maximise the prediction of trait variance. Genetic influences operate via the environment by necessity, therefore the pathways from genotype to phenotype operate through rGE mechanisms. Although within-family genetic effects do not include prGE effects due to between-family factors such as SES, within-family genetic effects are not free of *all* kinds of rGE, as demonstrated by twin studies showing that correlations between putative measures of the environment and children’s specific outcomes are genetically influenced^58^. Within-family GPS prediction estimates can be interpreted as direct genetic effects in the sense that they stem from the individual level not the family level. Children select, modify and create experiences (active rGE), or evoke responses in their environment (evocative rGE) that are correlated with their genetic propensities. Therefore, within-family genetic differences can relate to trait differences through active or evocative rGE pathways, but are free of any passive rGE effects.

### Implications

The results from this study have three important implications for the interpretation of the existing polygenic score literature, as well as for future genetic research. First, the finding that between-family predictions pick up effects due to prGE only in cognitive, and not non-cognitive traits is informative for causal inference studies that use designs such as Mendelian Randomisation^61,62^. Here, a genetic instrument that is related to a predictor (in form of a single genetic marker or GPS) is used to assess the causal relationship between the predictor and an outcome. At a population level, genotypes are not inherited randomly: individuals with particular genotypes are not born into environmental conditions at chance. If family environment is associated with the genetic instrument as well as the predictor and the outcome, this opens a backdoor path whereby predictor and outcome are related through the prGE mechanisms^19^. This could lead to an assumption violation, therefore biasing causal inference in between-family analysis. Only in a within-family design is it ensured that Mendelian Randomisation meets its assumptions because transmission of alleles is randomised at meiosis within families, and because prGE effects due to shared environment are held constant^27,63–65^. Although genetic data for siblings is often not available, our results provide a useful guideline for the GPS-outcome combinations that are unlikely to suffer from this assumption violation when applying designs such as Mendelian Randomisation to unrelated samples. For example, our results indicate that caution should be warranted due to prGE mechanisms if applying Mendelian Randomisation to cognitive traits, even if family SES is included as a confounder in the analyses as confounding effects might not be captured perfectly. On the contrary, other traits such as BMI and ADHD (with the possible exclusion of the self-rated health GPS) should be less problematic, because within- and between-family effect sizes match closely, ruling out potential confounding due to prGE.

Second, our results provide evidence that location-related population stratification is not a systematic bias in GPS prediction of complex traits when controlling for genetic principal components in samples from White European backgrounds. For example, for height and BMI, the decomposition of GPS effects showed that between-family prediction estimates were similar to within-family estimates, which are by necessity free of population stratification since stratification is constant within a family. For those traits where within- and between-family estimate differences were large and significant, differences disappeared after accounting for SES, indicating that SES was the main source of the discrepancy, as opposed to location-related population stratification.

Third, our study illustrates the usefulness of obtaining genotypic data on family members, since it makes it possible to identify mechanisms of polygenic prediction. Our results demonstrate that by analysing DZ co-twins’ genetic data jointly, prGE mechanisms due to shared environment (and in this case associated with SES) can be revealed.

### Limitations

Although we present the most comprehensive within- and between-family comparison of GPS prediction to date, there are limitations to this study. The GWA studies used to generate the eight GPS for this study had different statistical power to discover genetic effect sizes due to sample size variations and different underlying genetic architectures of the GWA study traits. As a result, each of the eight GPS were differently powered to detect target- and cross-trait associations, making it difficult to draw direct comparisons across the within- and between-family prediction effect sizes. Lack of power may also lead to an inability to detect small prGE effects that would become visible with (i) more powerful GPS and (ii) the availability of larger DZ twin pair samples. However, we detected prGE effects in cross-trait analysis using the ADHD GPS, which is based on the smallest GWAS study sample (~55,000 individuals), indicating that we had sufficient power to detect at least some of the prGE effects.

Another limitation was that we did not have parental genotypes available to directly test the influence of non-transmitted parental alleles on offspring outcomes (genetic nurture)^20^. Although the within-family design used in this study accounts for the effects of both transmitted and non-transmitted parental alleles on offspring outcomes, it is not possible to disentangle these two sources of prGE. Future studies would benefit from incorporating parental and sibling genotypes to disentangle the prGE effects through the joint analysis of parental and sibling genotypes, which will shed light on how both non-transmitted parental and non-co-inherited sibling alleles contribute to trait development.

### Conclusion

This study provided strong evidence for prGE mechanisms in polygenic score prediction for cognitive, but not non-cognitive, traits across a range of different polygenic scores. The implications of these findings for future studies depend on their aims. If maximising trait prediction is the goal, the use of unrelated samples is valid even in the presence of prGE effects because these influences are informative nonetheless. However, if the goal is causal inference and explanation, a within-family genetic design is recommended to avoid prGE-related confounding. The increasing availability of genotypic data in relatives will become a crucial element in genetics research, allowing researchers to disentangle the mechanisms of polygenic prediction of complex human traits.

## Online Resources

OSF pre-registration link

https://osf.io/eq8ga/?view_only=768f42366d134ebeb50f5999763c3fce

## Acknowledgements

We gratefully acknowledge the ongoing contribution of the participants in the Twins Early Development Study (TEDS) and their families. TEDS is supported by a programme grant to RP from the UK Medical Research Council (MR/M021475/1 and previously G0901245), with additional support from the US National Institutes of Health (AG046938). The research leading to these results has also received funding from the European Research Council under the European Union’s Seventh Framework Programme (FP7/2007-2013)/ grant agreement n° 602768 and ERC grant agreement n° 295366. RP is supported by a Medical Research Council Professorship award (G19/2). SS is supported by the MRC/IoPPN Excellence Award and by the US National Institutes of Health (AG046938). PFO received funding from the UK Medical Research Council (MR/N015746/1). High performance computing facilities were funded with capital equipment grants from the GSTT Charity (TR130505) and Maudsley Charity (980).

## Author Contributions

Study concept and design: S.S., R.P. Genotype quality control and polygenic score calculation: S.S. Analysis of data: S.S. Interpretation of data: S.S., R.P. Wrote the manuscript: S.S. Contributed and critically reviewed the manuscript: All authors

## References

1. Gratten, J., Wray, N.R., Keller, M.C., and Visscher, P.M. (2014). Large-scale genomics unveils the genetic architecture of psychiatric disorders. Nature Neuroscience 17, 782–790.

2. Plomin, R., and Stumm von, S. (2018). The new genetics of intelligence. Nat. Rev. Genet. 19, 148–159.

3. Martin, A.R., Daly, M.J., Robinson, E.B., Hyman, S.E., and Neale, B.M. (2018). Predicting polygenic risk of psychiatric disorders. Biological Psychiatry.

4. Wray, N.R., Lee, S.H., Mehta, D., Vinkhuyzen, A.A.E., Dudbridge, F., and Middeldorp, C.M. (2014). Research review: Polygenic methods and their application to psychiatric traits. J Child Psychol Psychiatry 55, 1068–1087.

5. Plomin, R. (2018). Blueprint (Penguin UK).

6. Lee, J.J., Wedow, R., Okbay, A., Kong, E., Maghzian, O., Zacher, M., Nguyen-Viet, T.A., Bowers, P., Sidorenko, J., Linner, R.K., et al. (2018). Gene discovery and polygenic prediction from a genome-wide association study of educational attainment in 1.1 million individuals. Nat. Genet. 50, 1112–1121.

7. Allegrini, A., Selzam, S., Rimfeld, K., Stumm von, S., Pingault, J.-B., and Plomin, R. (2019). Genomic prediction of cognitive traits in childhood and adolescence. Molecular Psychiatry.

8. Hill, W.D., Hagenaars, S.P., Marioni, R.E., Harris, S.E., Liewald, D.C.M., Davies, G., Okbay, A., McIntosh, A.M., Gale, C.R., and Deary, I.J. (2016). Molecular Genetic Contributions to Social Deprivation and Household Income in UK Biobank. Curr. Biol. 26, 3083–3089.

9. Selzam, S., Krapohl, E., Stumm von S., O’Reilly, P.F., Rimfeld, K., Kovas, Y., Dale, P.S., Lee, J.J., and Plomin, R. (2017). Predicting educational achievement from DNA. Molecular Psychiatry 22, 267–272.

10. Belsky, D.W., Domingue, B.W., Wedow, R., Arseneault, L., Boardman, J.D., Caspi, A., Conley, D., Fletcher, J.M., Freese, J., Herd, P., et al. (2018). Genetic analysis of social-class mobility in five longitudinal studies. Proc. Natl. Acad. Sci. U.S.a. 115, E7275–E7284.

11. Belsky, D.W., Moffitt, T.E., Corcoran, D.L., Domingue, B., Harrington, H., Hogan, S., Houts, R., Ramrakha, S., Sugden, K., Williams, B.S., et al. (2016). The Genetics of Success. Psychological Science 27, 957–972.

12. de Zeeuw, E.L., van Beijsterveldt, C.E.M., Glasner, T.J., Bartels, M., Ehli, E.A., Davies, G.E., Hudziak, J.J., Rietveld, C.A., Groen-Blokhuis, M.M., Hottenga, J.-J., et al. (2014). Polygenic Scores Associated With Educational Attainment in Adults Predict Educational Achievement and ADHD Symptoms in Children. Am. J. Med. Genet. B Neuropsychiatr. Genet. 165, 510–520.

13. Hagenaars, S.P., Harris, S.E., Davies, G., Hill, W.D., Liewald, D.C.M., Ritchie, S.J., Marioni, R.E., Fawns-Ritchie, C., Cullen, B., Malik, R., et al. (2016). Shared genetic aetiology between cognitive functions and physical and mental health in UK Biobank (N=112 151) and 24 GWAS consortia. Molecular Psychiatry 21, 1624–1632.

14. Mõttus, R., Realo, A., Vainik, U., Allik, J., and Esko, T. (2017). Educational Attainment and Personality Are Genetically Intertwined. Psychological Science 28, 1631–1639.

15. Smith-Woolley, E., Selzam, S., and Plomin, R. (2019). Polygenic score for educational attainment captures DNA variants shared between personality traits and educational achievement. J Pers Soc Psychol.

16. Krapohl, E., Euesden, J., Zabaneh, D., Pingault, J.-B., Rimfeld, K., Stumm von S., Dale, P.S., Breen, G., O’Reilly, P.F., and Plomin, R. (2016). Phenome-wide analysis of genome-wide polygenic scores. Molecular Psychiatry 21, 1188–1193.

17. Belsky, D.W., and Harden, K.P. (2019). Phenotypic Annotation: Using Polygenic Scores to Translate Discoveries From Genome-Wide Association Studies From the Top Down. Current Directions in Psychological Science 096372141880772.

18. Plomin, R., DeFries, J.C., and Loehlin, J.C. (1977). Genotype-environment interaction and correlation in the analysis of human behavior. Psychol Bull 84, 309–322.

19. Pingault, J.-B., O’Reilly, P.F., Schoeler, T., Ploubidis, G.B., Rijsdijk, F., and Dudbridge, F. (2018). Using genetic data to strengthen causal inference in observational research. Nat. Rev. Genet. 19, 566–580.

20. Kong, A., Thorleifsson, G., Frigge, M.L., Vilhjalmsson, B.J., Young, A.I., Thorgeirsson, T.E., Benonisdottir, S., Oddsson, A., Halldorsson, B.V., Masson, G., et al. (2018). The nature of nurture: Effects of parental genotypes. Science 359, 424–428.

21. Bates, T.C., Maher, B.S., Medland, S.E., McAloney, K., Wright, M.J., Hansell, N.K., Kendler, K.S., Martin, N.G., and Gillespie, N.A. (2018). The Nature of Nurture: Using a Virtual-Parent Design to Test Parenting Effects on Children’s Educational Attainment in Genotyped Families. Twin Res Hum Genet. 21, 73–83.

22. Koellinger, P.D., and Harden, K.P. (2018). Using nature to understand nurture. Science 359, 386–387.

23. Fisher, R.A. (1918). The Correlation between Relatives on the Supposition of Mendelian Inheritance. Transactions of the Royal Society of Edinburgh 52, 399–433.

24. Fletcher, J.M. (2011). The promise and pitfalls of combining genetic and economic research. Health Economics 20, 889–892.

25. Young, A.I., Frigge, M.L., Gudbjartsson, D.F., Thorleifsson, G., Bjornsdottir, G., Sulem, P., Masson, G., Thorsteinsdottir, U., Stefansson, K., and Kong, A. (2018). Relatedness disequilibrium regression estimates heritability without environmental bias. Nat. Genet. 50, 1304–1310.

26. Benyamin, B., Visscher, P.M., and McRae, A.F. (2009). Family-based genome-wide association studies. Pharmacogenomics 10, 181–190.

27. Brumpton, B., Sanderson, E., Pires Hartwig, F., Harrison, S., Aberge Vie, G., Cho, Y., Hughes, A., Boomsma, D., Havdahl, A., Hopper, J., et al. (2019). Within-family studies for Mendelian randomization: avoiding dynastic, assortative mating, and population stratification biases. bioRxiv.

28. Domingue, B.W., Belsky, D.W., Conley, D., Harris, K.M., and Boardman, J.D. (2015). Polygenic Influence on Educational Attainment. AERA Open 1, 1–13.

29. Rietveld, C.A., Esko, T., Davies, G., Pers, T.H., Turley, P., Benyamin, B., Chabris, C.F., Emilsson, V., Johnson, A.D., Lee, J.J., et al. (2014). Common genetic variants associated with cognitive performance identified using the proxy-phenotype method. Proc. Natl. Acad. Sci. U.S.a. 111, 13790–13794.

30. Spinath, B., Spinath, F.M., Harlaar, N., and Plomin, R. (2006). Predicting school achievement from general cognitive ability, self-perceived ability, and intrinsic value. Intelligence 34, 363–374.

31. Krapohl, E., Rimfeld, K., Shakeshaft, N.G., Trzaskowski, M., McMillan, A., Pingault, J.-B., Asbury, K., Harlaar, N., Kovas, Y., Dale, P.S., et al. (2014). The high heritability of educational achievement reflects many genetically influenced traits, not just intelligence. Proc. Natl. Acad. Sci. U.S.a. 111, 15273–15278.

32. Briley, D.A., Domiteaux, M., and Tucker-Drob, E.M. (2014). Achievement-Relevant Personality: Relations with the Big Five and Validation of an Efficient Instrument. Learning and Individual Differences 32, 26–39.

33. Marques, S.C., Pais-Ribeiro, J.L., and Lopez, S.J. (2011). The Role of Positive Psychology Constructs in Predicting Mental Health and Academic Achievement in Children and Adolescents: A Two-Year Longitudinal Study. J Happiness Stud 12, 1049–1062.

34. Pingault, J.-B., Tremblay, R.E., Vitaro, F., Carbonneau, R., Genolini, C., Falissard, B., and Cote, S.M. (2011). Childhood Trajectories of Inattention and Hyperactivity and Prediction of Educational Attainment in Early Adulthood: A 16-Year Longitudinal Population-Based Study. American Journal of Psychiatry 168, 1164–1170.

35. De Ridder, K.A.A., Pape, K., Johnsen, R., Holmen, T.L., Westin, S., and Bjorngaard, J.H. (2013). Adolescent Health and High School Dropout: A Prospective Cohort Study of 9000 Norwegian Adolescents (The Young-HUNT). Plos On 8.

36. Zuffianò, A., Alessandri, G., Gerbino, M., Luengo Kanacri, B.P., Di Giunta, L., Milioni, M., and Caprara, G.V. (2013). Academic achievement: The unique contribution of self-efficacy beliefs in self-regulated learning beyond intelligence, personality traits, and self-esteem. Learning and Individual Differences 23, 158–162.

37. Krapohl, E., and Plomin, R. (2016). Genetic link between family socioeconomic status and children’s educational achievement estimated from genome-wide SNPs. Molecular Psychiatry 21, 437–443.

38. Trzaskowski, M., Harlaar, N., Arden, R., Krapohl, E., Rimfeld, K., McMillan, A., Dale, P.S., and Plomin, R. (2014). Genetic influence on family socioeconomic status and children’s intelligence. Intelligence 42, 83–88.

39. Haworth, C.M.A., Davis, O.S.P., and Plomin, R. (2013). Twins Early Development Study (TEDS): A Genetically Sensitive Investigation of Cognitive and Behavioral Development From Childhood to Young Adulthood. Twin Res Hum Genet. 16, 117–125.

40. Ware, J.E., and Sherbourne, C.D. (1992). The MOS 36-item short-form health survey (SF-36). I. Conceptual framework and item selection. Med Care 30, 473–483.

41. Raven, J., and Court, J.H. (1996). Manual for Raven’s Progressive Matrices and Vocabulary Scales (Oxford: Oxford University Press).

42. Wechsler, D. (1992). Wechsler Intelligence Scale for Children - Third Edition UK (WISC-III- UK) Manual (London: The Psychological Corporation).

43. Kaplan, E., Fein, D., Kramer, J., Delis, D., and Morris, R. (1999). WISC-III as a Process Instrument (WISC-III-PI) (New York: The Psychological Corporation).

44. Mullins-Sweatt, S.N., Jamerson, J.E., Samuel, D.B., Olson, D.R., and Widiger, T.A. (2006). Psychometric properties of an abbreviated instrument of the five-factor model. Assessment 13, 119–137.

45. McInnes, G., Tanigawa, Y., DeBoever, C., Lavertu, A., Olivieri, J.E., Aguirre, M., and Rivas, M.A. (2018). Global Biobank Engine: enabling genotype-phenotype browsing for biobank summary statistics. Bioinformatics 9, 1612.

46. Harris, S.E., Hagenaars, S.P., Davies, G., Hill, W.D., Liewald, D.C.M., Ritchie, S.J., Marioni, R.E., Sudlow, C.L.M., Wardlaw, J.M., McIntosh, A.M., et al. (2017). Molecular genetic contributions to self-rated health. Int J Epidemiol 46, 994–1009.

47. Savage, J.E., Jansen, P.R., Stringer, S., Watanabe, K., Bryois, J., de Leeuw, C.A., Nagel, M., Awasthi, S., Barr, P.B., Coleman, J.R.I., et al. (2018). Genome-wide association meta-analysis in 269,867 individuals identifies new genetic and functional links to intelligence. Nat. Genet. 50, 912–919.

48. Yengo, L., Sidorenko, J., Kemper, K.E., Zheng, Z., Wood, A.R., Weedon, M.N., Frayling, T.M., Hirschhorn, J., Yang, J., Visscher, P.M., et al. (2018). Meta-analysis of genome-wide association studies for height and body mass index in similar to 700 000 individuals of European ancestry. Hum. Mol. Genet. 27, 3641–3649.

49. Luciano, M., Hagenaars, S.P., Davies, G., Hill, W.D., Clarke, T.-K., Shirali, M., Harris, S.E., Marioni, R.E., Liewald, D.C., Fawns-Ritchie, C., et al. (2018). Association analysis in over 329,000 individuals identifies 116 independent variants influencing neuroticism. Nat. Genet. 50, 6–11.

50. Demontis, D., Walters, R.K., Martin, J., Mattheisen, M., Als, T.D., Agerbo, E., Baldursson, G., Belliveau, R., Bybjerg-Grauholm, J., Bækvad-Hansen, M., et al. (2019). Discovery of the first genome-wide significant risk loci for attention deficit/hyperactivity disorder. Nat. Genet. 51, 63–75.

51. Pardiñas, A.F., Holmans, P., Pocklington, A.J., Escott-Price, V., Ripke, S., Carrera, N., Legge, S.E., Bishop, S., Cameron, D., Hamshere, M.L., et al. (2018). Common schizophrenia alleles are enriched in mutation-intolerant genes and in regions under strong background selection. Nat. Genet. 50, 381–389.

52. Vilhjalmsson, B.J., Yang, J., Finucane, H.K., Gusev, A., Lindstrom, S., Ripke, S., Genovese, G., Loh, P.-R., Bhatia, G., Do, R., et al. (2015). Modeling Linkage Disequilibrium Increases Accuracy of Polygenic Risk Scores. Am. J. Hum. Genet. 97, 576–592.

53. Snijders, T.A.B., and Bosker, R.J. (1999). Multilevel Analysis (SAGE).

54. Carlin, J.B., Gurrin, L.C., Sterne, J., Morley, R., and Dwyer, T. (2005). Regression models for twin studies: a critical review. Int J Epidemiol 34, 1089–1099.

55. Genser, B., Teles, C.A., Barreto, M.L., and Fischer, J.E. (2015). Within- and between- group regression for improving the robustness of causal claims in cross-sectional analysis. Environmental Health 2015 14:1 14, 60.

56. North, BV, Curtis, D., and Sham, P.C. (2002). A note on the calculation of empirical P values from Monte Carlo procedures. The American Journal of Human Genetics 71, 439–441.

57. Benjamini, Y., and Hochberg, Y. (1995). Controlling the False Discovery Rate: a Practical and Powerful Approach to Multiple Testing. Journal of the Royal Statistical Society. Series B (Statistical Methodology) 57, 289–300.

58. Plomin, R. (1994). Genetics and experience: The interplay between nature and nurture. (Sage Publications, Inc).

59. Polderman, T.J.C., Benyamin, B., de Leeuw, C.A., Sullivan, P.F., van Bochoven, A., Visscher, P.M., and Posthuma, D. (2015). Meta-analysis of the heritability of human traits based on fifty years of twin studies. Nat. Genet. 47, 702–709.

60. Rijsdijk, F.V. (2002). Analytic approaches to twin data using structural equation models. Briefings in Bioinformatics 3, 119–133.

61. Smith, G.D., and Ebrahim, S. (2005). What can mendelian randomisation tell us about modifiable behavioural and environmental exposures? Bmj 330, 1076–1079.

62. Smith, G.D., and Hemani, G. (2014). Mendelian randomization: genetic anchors for causal inference in epidemiological studies. Hum. Mol. Genet. 23, R89–R98.

63. Pingault, J.-B., O’Reilly, P.F., Schoeler, T., Ploubidis, G.B., Rijsdijk, F., and Dudbridge, F. (2018). Using genetic data to strengthen causal inference in observational research. Nat. Rev. Genet. 19, 566–580.

64. Smith, G.D., and Ebrahim, S. (2003). “Mendelian randomization”: can genetic epidemiology contribute to understanding environmental determinants of disease? Int J Epidemiol 32, 1–22.

65. Smith, G.D. (2007). Capitalizing on Mendelian randomization to assess the effects of treatments. J R Soc Med 100, 432–435.

