## Supplementary Materials for "Comparing within- and between-family polygenic score prediction"

#### Supplementary Online Material

### Supplementary Tables

Table S1. Descriptive statistics, age and sex effects for phenotypes

| | N Pairs | Mean | SD | Skew | Min | Max | F sex | P sex | $R^2$ sex | F age | P age | $R^2$ age | P order | $R^2$ order |
|---|---|---|---|---|---|---|---|---|---|---|---|---|---|---|
| **Height** | 1,463 | 171.997 | 10.457 | 0.142 | 132 | 211 | 1,460.639 | < 0.001 | 0.500 | 0.855 | 0.355 | 0.001 | 0.174 | <0.001 |
| **BMI** | 1,353 | 23.495 | 4.676 | 1.520 | 12.061 | 47.477 | 0.002 | 0.964 | <0.001 | 16.470 | < 0.001 | 0.012 | 0.918 | <0.001 |
| **Self-rated Health** | 1,494 | 3.480 | 0.672 | -0.330 | 1.000 | 5.000 | 8.188 | 0.004 | 0.005 | 1.457 | 0.228 | 0.001 | 0.987 | <0.001 |
| **IQ** | 1,569 | 0.117 | 0.954 | -0.242 | -3.441 | 3.040 | 7.277 | 0.007 | 0.005 | 104.278 | < 0.001 | 0.062 | 0.125 | <0.001 |
| **GCSE** | 2,366 | 8.952 | 1.194 | -0.330 | 4.670 | 11.000 | 1.814 | 0.178 | <0.001 | 4.49 | 0.034 | 0.001 | 0.487 | <0.001 |
| **Neuroticism** | 789 | 2.583 | 0.655 | 0.280 | 1.000 | 5.000 | 31.894 | < 0.001 | 0.039 | 2.202 | 0.138 | 0.003 | 0.740 | <0.001 |
| **ADHD Symptoms** | 2,469 | 0.063 | 1.002 | 1.357 | -1.371 | 5.066 | 159.896 | < 0.001 | 0.061 | 16.877 | < 0.001 | 0.007 | 0.073 | <0.001 |
| **SCZ Symptoms** | 1,140 | -0.026 | 0.705 | 1.560 | -0.816 | 4.093 | 2.041 | 0.153 | 0.002 | 6.122 | 0.013 | 0.005 | 0.858 | <0.001 |
| **SES** | 2,962 | 0.209 | 0.994 | 0.046 | -2.351 | 2.495 | -- | -- | -- | -- | -- | -- | -- | -- |

**Note.** Means and standard deviations for individual measures are calculated based on raw data. Height, BMI, self-reported health, GCSE grades and neuroticism means and standard deviations are reported on their original scale. IQ, ADHD symptoms, schizophrenia symptoms and socioeconomic status are reported on the z-scale as standardization was required to form the composite. Sex, age and birth order tests were performed on one randomly selected twin per pair. $R^2$= proportion of variance explained. Order = birth order; BMI = Body Mass Index; IQ = Intelligence; GCSE = General Certificate of Secondary Education (educational achievement); ADHD = Attention-Deficit/Hyperactivity Disorder; SCZ = Schizophrenia; SES = family socio-economic status.

Table S2. GWAS used for polygenic score calculation

| Trait | Year | SNP-$h^2$ | GPS $R^2$ | Cases | Controls | GWAS sample size | Notes |
|---|---|---|---|---|---|---|---|
| BMI[1] | 2018 | 22.4% (3.7%)[1] | 10.2%[1] | - | - | 681,275 | - |
| Height[1] | 2018 | 48.3% (3.7%)[1] | 24.4%[1] | - | - | 693,529 | - |
| Self-rated health[2] | 2018 | 13% (0. 6%)[3] | -- | - | - | 337,199 | - |
| Intelligence[4] | 2018 | 19% (1%)[4] | 6.7%[5] | - | - | 266,453 | GWAS re-run excluding TEDS sample (3,414) |
| Educational Attainment[6] | 2018 | 12.2% (0.3%)[6] | 11.4% [6] | - | - | 766,345 | - |
| Neuroticism[7] | 2017 | 10.8% (0.5%)[7] | 2.8% [7] | - | - | 329,821 | - |
| ADHD[8] | 2019 | 21.6% (1.4%)[8] | 5.5%[8] | 20,183 | 35,191 | 55,374 | - |
| Schizophrenia[9] | 2018 | 20% (0.6%)[9] | 5.7%[9] | 40,675 | 64,643 | 105,318 | - |

**Note.** $H^2$ = heritability; $R^2$ = phenotypic variance explained.

Table S3. Intraclass coefficients and residual effects

| Phenotype | N pairs | ICC | ICC 95% CI lower | ICC 95% CI upper | RandEff.resid |
|---|---|---|---|---|---|
| **Height** | 1,463 | 0.439 | 0.386 | 0.497 | 0.561 |
| **BMI** | 1,353 | 0.317 | 0.266 | 0.377 | 0.683 |
| **Self-rated Health** | 1,494 | 0.136 | 0.095 | 0.196 | 0.863 |
| **IQ** | 1,569 | 0.422 | 0.372 | 0.478 | 0.578 |
| **GCSE** | 2,366 | 0.582 | 0.537 | 0.63 | 0.418 |
| **Neuroticism** | 789 | 0.103 | 0.052 | 0.203 | 0.897 |
| **ADHD Symptoms** | 2,469 | 0.323 | 0.284 | 0.367 | 0.677 |
| **SCZ Symptoms** | 1,140 | 0.254 | 0.201 | 0.32 | 0.746 |

**Note.** BMI = Body Mass Index; IQ = Intelligence; GCSE = General Certificate of Secondary Education (educational achievement); ADHD = Attention-Deficit/Hyperactivity Disorder; SCZ = Schizophrenia symptoms; ICC = Intraclass coefficient; CI = Confidence Interval; RandEff.Resid = Residual of the random effect.

Table S4. Within- and between-family prediction estimates

| Phenotype | GPS | beta.B | SE.B | L.CI.B | U.CI.B | P.B | beta.W | SE.W | CI.L.W | CI.U.W | P.W | TotEff | PercReduc | P.diff |
|---|---|---|---|---|---|---|---|---|---|---|---|---|---|---|
| ADHD | ADHD | 0.111 | 0.019 | 0.085 | 0.137 | 6.80e-09 | 0.124 | 0.023 | 0.072 | 0.176 | 1.50e-07 | 0.120 | 11.530 | 0.681 |
| ADHD | BMI | 0.093 | 0.019 | 0.067 | 0.120 | 6.92e-07 | 0.038 | 0.024 | -0.014 | 0.088 | 0.114 | 0.056 | 59.670 | 0.034 |
| ADHD | EA | -0.118 | 0.018 | -0.143 | -0.093 | 1.40e-10 | -0.140 | 0.025 | -0.197 | -0.083 | 3.09e-08 | -0.133 | 18.660 | 0.38 |
| ADHD | Height | -0.020 | 0.019 | -0.045 | 0.006 | 0.288 | 0.017 | 0.024 | -0.032 | 0.068 | 0.464 | 0.005 | 187.390 | 0.252 |
| ADHD | IQ | -0.106 | 0.019 | -0.131 | -0.081 | 1.22e-08 | -0.122 | 0.024 | -0.173 | -0.071 | 6.38e-07 | -0.117 | 14.640 | 0.568 |
| ADHD | Neurot | 0.044 | 0.019 | 0.018 | 0.071 | 0.021 | 0.052 | 0.023 | 0.003 | 0.102 | 0.023 | 0.050 | 18.170 | 0.924 |
| ADHD | SCZ | -0.001 | 0.019 | -0.026 | 0.024 | 0.963 | 0.039 | 0.024 | -0.012 | 0.090 | 0.107 | 0.026 | 4,500.790 | 0.127 |
| ADHD | SRH | -0.125 | 0.019 | -0.152 | -0.099 | 2.78e-11 | -0.035 | 0.025 | -0.088 | 0.019 | 0.151 | -0.064 | 71.720 | 3.90e-04 |
| BMI | ADHD | 0.068 | 0.026 | 0.033 | 0.102 | 0.008 | 0.059 | 0.032 | -0.009 | 0.128 | 0.064 | 0.062 | 12.720 | 0.719 |
| BMI | BMI | 0.350 | 0.024 | 0.316 | 0.383 | 2.35e-46 | 0.297 | 0.031 | 0.233 | 0.361 | 1.76e-21 | 0.314 | 15.110 | 0.05 |
| BMI | EA | -0.113 | 0.025 | -0.149 | -0.078 | 9.69e-06 | -0.066 | 0.034 | -0.141 | 0.009 | 0.053 | -0.081 | 40.950 | 0.13 |
| BMI | Height | -0.047 | 0.025 | -0.081 | -0.013 | 0.056 | -0.032 | 0.032 | -0.102 | 0.041 | 0.319 | -0.037 | 32.930 | 0.675 |
| BMI | IQ | -0.048 | 0.025 | -0.082 | -0.013 | 0.058 | 0.010 | 0.033 | -0.064 | 0.083 | 0.76 | -0.008 | 120.870 | 0.153 |
| BMI | Neurot | -0.081 | 0.026 | -0.116 | -0.046 | 0.002 | -0.007 | 0.031 | -0.071 | 0.058 | 0.821 | -0.030 | 91.220 | 0.12 |
| BMI | SCZ | -0.050 | 0.026 | -0.085 | -0.015 | 0.049 | -0.055 | 0.032 | -0.125 | 0.012 | 0.085 | -0.054 | 9.630 | 0.526 |
| BMI | SRH | -0.143 | 0.026 | -0.180 | -0.107 | 3.20e-08 | -0.133 | 0.032 | -0.200 | -0.068 | 3.56e-05 | -0.136 | 6.620 | 0.96 |
| GCSE | ADHD | -0.176 | 0.021 | -0.200 | -0.153 | 7.32e-17 | -0.061 | 0.019 | -0.105 | -0.017 | 0.001 | -0.128 | 65.370 | 2.00e-05 |
| GCSE | BMI | -0.135 | 0.021 | -0.159 | -0.112 | 1.88e-10 | -0.023 | 0.019 | -0.068 | 0.023 | 0.239 | -0.088 | 83.250 | 9.99e-06 |
| GCSE | EA | 0.418 | 0.019 | 0.397 | 0.440 | 1.60e-98 | 0.214 | 0.020 | 0.167 | 0.260 | 2.22e-26 | 0.333 | 48.930 | 9.99e-06 |
| GCSE | Height | 0.033 | 0.021 | 0.009 | 0.057 | 0.108 | 0 | 0.019 | -0.046 | 0.046 | 0.982 | 0.019 | 101.310 | 0.163 |
| GCSE | IQ | 0.320 | 0.020 | 0.298 | 0.342 | 6.21e-55 | 0.201 | 0.019 | 0.156 | 0.247 | 7.24e-25 | 0.270 | 37.210 | 9.99e-06 |
| GCSE | Neurot | -0.074 | 0.021 | -0.099 | -0.051 | 5.26e-04 | -0.061 | 0.018 | -0.103 | -0.018 | 8.86e-04 | -0.069 | 18.400 | 0.472 |
| GCSE | SCZ | 0.039 | 0.021 | 0.016 | 0.063 | 0.063 | -0.017 | 0.019 | -0.063 | 0.030 | 0.378 | 0.016 | 143.240 | 0.025 |

| | | | | | | | | | | | | | | |
|---|---|---|---|---|---|---|---|---|---|---|---|---|---|---|
| GCSE | SRH | 0.188 | 0.021 | 0.165 | 0.211 | 4.93e-19 | 0.057 | 0.019 | 0.012 | 0.103 | 0.003 | 0.133 | 69.510 | 9.99e-06 |
| Height | ADHD | -0.035 | 0.026 | -0.068 | -0.002 | 0.176 | -0.057 | 0.028 | -0.121 | 0.008 | 0.042 | -0.047 | 60.850 | 0.26 |
| Height | BMI | -0.021 | 0.026 | -0.054 | 0.011 | 0.414 | 0.025 | 0.028 | -0.037 | 0.087 | 0.368 | 0.005 | 219.220 | 0.252 |
| Height | EA | 0.034 | 0.026 | 0.002 | 0.067 | 0.181 | -0.004 | 0.030 | -0.074 | 0.064 | 0.888 | 0.013 | 112.350 | 0.168 |
| Height | Height | 0.465 | 0.022 | 0.438 | 0.493 | 7.48e-90 | 0.410 | 0.026 | 0.350 | 0.470 | 5.72e-53 | 0.435 | 11.830 | 0.086 |
| Height | IQ | 0.042 | 0.025 | 0.009 | 0.074 | 0.098 | -0.006 | 0.029 | -0.071 | 0.057 | 0.825 | 0.015 | 115.250 | 0.124 |
| Height | Neurot | -0.058 | 0.026 | -0.091 | -0.025 | 0.025 | 0.007 | 0.027 | -0.055 | 0.071 | 0.799 | -0.022 | 112.010 | 0.081 |
| Height | SCZ | -0.051 | 0.026 | -0.084 | -0.018 | 0.046 | 0.015 | 0.028 | -0.045 | 0.078 | 0.583 | -0.014 | 130.070 | 0.107 |
| Height | SRH | 0.057 | 0.026 | 0.025 | 0.088 | 0.027 | -0.027 | 0.028 | -0.087 | 0.032 | 0.344 | 0.010 | 146.960 | 0.126 |
| IQ | ADHD | -0.085 | 0.024 | -0.117 | -0.054 | 4.48e-04 | -0.004 | 0.027 | -0.066 | 0.057 | 0.874 | -0.039 | 94.920 | 0.012 |
| IQ | BMI | -0.029 | 0.024 | -0.058 | 0.002 | 0.241 | 0.014 | 0.027 | -0.047 | 0.076 | 0.6 | -0.004 | 149.710 | 0.264 |
| IQ | EA | 0.253 | 0.023 | 0.224 | 0.283 | 1.12e-26 | 0.126 | 0.030 | 0.061 | 0.192 | 2.15e-05 | 0.180 | 50.090 | 6.00e-05 |
| IQ | Height | 0.009 | 0.024 | -0.022 | 0.040 | 0.716 | 0.044 | 0.027 | -0.022 | 0.110 | 0.099 | 0.029 | 399.060 | 0.562 |
| IQ | IQ | 0.263 | 0.024 | 0.233 | 0.292 | 2.86e-27 | 0.137 | 0.028 | 0.074 | 0.196 | 1.32e-06 | 0.190 | 47.980 | 3.50e-04 |
| IQ | Neurot | -0.018 | 0.025 | -0.050 | 0.013 | 0.457 | -0.018 | 0.026 | -0.079 | 0.043 | 0.496 | -0.018 | 2 | 0.686 |
| IQ | SCZ | 0.006 | 0.025 | -0.025 | 0.037 | 0.824 | 0.012 | 0.027 | -0.048 | 0.074 | 0.653 | 0.009 | 122.300 | 0.408 |
| IQ | SRH | 0.109 | 0.024 | 0.078 | 0.139 | 8.19e-06 | 0.022 | 0.029 | -0.043 | 0.087 | 0.435 | 0.059 | 79.420 | 0.008 |
| Neurot | ADHD | -0.009 | 0.031 | -0.059 | 0.039 | 0.765 | 0.081 | 0.048 | -0.010 | 0.172 | 0.092 | 0.071 | 963.990 | 0.341 |
| Neurot | BMI | -0.017 | 0.031 | -0.067 | 0.035 | 0.586 | 0.020 | 0.047 | -0.076 | 0.116 | 0.662 | 0.017 | 220.620 | 0.302 |
| Neurot | EA | 0.015 | 0.029 | -0.032 | 0.061 | 0.601 | 0.015 | 0.054 | -0.093 | 0.129 | 0.779 | 0.015 | 0.220 | 0.947 |
| Neurot | Height | -0.025 | 0.030 | -0.074 | 0.024 | 0.407 | 0.036 | 0.047 | -0.057 | 0.127 | 0.454 | 0.029 | 242.680 | 0.245 |
| Neurot | IQ | -0.006 | 0.030 | -0.055 | 0.041 | 0.849 | 0.057 | 0.050 | -0.044 | 0.161 | 0.249 | 0.051 | 1,085.060 | 0.203 |
| Neurot | Neurot | 0.108 | 0.032 | 0.055 | 0.160 | 6.34e-04 | 0.021 | 0.047 | -0.076 | 0.117 | 0.649 | 0.030 | 80.410 | 0.068 |
| Neurot | SCZ | -0.014 | 0.031 | -0.061 | 0.034 | 0.662 | 0.063 | 0.049 | -0.035 | 0.162 | 0.192 | 0.056 | 569.820 | 0.212 |
| Neurot | SRH | -0.041 | 0.030 | -0.088 | 0.006 | 0.176 | -0.024 | 0.049 | -0.124 | 0.074 | 0.617 | -0.026 | 39.940 | 0.499 |

| | | | | | | | | | | | | | | |
|---|---|---|---|---|---|---|---|---|---|---|---|---|---|---|
| SCZ | ADHD | 0.044 | 0.028 | 0.006 | 0.082 | 0.118 | 0.003 | 0.037 | -0.073 | 0.080 | 0.943 | 0.013 | 94.060 | 0.67 |
| SCZ | BMI | 0.109 | 0.027 | 0.070 | 0.146 | 5.72e-05 | 0.040 | 0.037 | -0.040 | 0.120 | 0.276 | 0.058 | 63.060 | 0.142 |
| SCZ | EA | -0.081 | 0.027 | -0.120 | -0.042 | 0.003 | 0.009 | 0.039 | -0.072 | 0.092 | 0.811 | -0.014 | 111.540 | 0.014 |
| SCZ | Height | 0 | 0.026 | -0.039 | 0.039 | 0.991 | 0.002 | 0.037 | -0.070 | 0.074 | 0.952 | 0.002 | 832.510 | 0.94 |
| SCZ | IQ | -0.024 | 0.027 | -0.062 | 0.015 | 0.366 | 0.025 | 0.038 | -0.054 | 0.106 | 0.51 | 0.013 | 204.800 | 0.333 |
| SCZ | Neurot | 0.062 | 0.028 | 0.025 | 0.098 | 0.027 | -0.024 | 0.035 | -0.093 | 0.048 | 0.501 | -0.002 | 138.190 | 0.071 |
| SCZ | SCZ | -0.010 | 0.027 | -0.048 | 0.029 | 0.715 | -0.011 | 0.037 | -0.092 | 0.068 | 0.777 | -0.010 | 4.950 | 0.854 |
| SCZ | SRH | -0.122 | 0.027 | -0.158 | -0.085 | 9.91e-06 | -0.012 | 0.037 | -0.089 | 0.066 | 0.741 | -0.040 | 90.060 | 0.015 |
| SRH | ADHD | -0.056 | 0.023 | -0.091 | -0.020 | 0.014 | -0.071 | 0.034 | -0.141 | -0.002 | 0.038 | -0.069 | 27.060 | 0.69 |
| SRH | BMI | -0.107 | 0.022 | -0.141 | -0.073 | 2.07e-06 | -0.011 | 0.034 | -0.080 | 0.055 | 0.752 | -0.024 | 89.870 | 0.091 |
| SRH | EA | 0.069 | 0.022 | 0.035 | 0.104 | 0.002 | 0.079 | 0.037 | 0.006 | 0.153 | 0.031 | 0.078 | 14.170 | 0.94 |
| SRH | Height | 0.062 | 0.022 | 0.028 | 0.096 | 0.004 | -0.008 | 0.034 | -0.077 | 0.060 | 0.806 | 0.001 | 113.560 | 0.054 |
| SRH | IQ | 0.022 | 0.022 | -0.013 | 0.057 | 0.316 | 0.027 | 0.035 | -0.049 | 0.101 | 0.445 | 0.026 | 20.820 | 0.365 |
| SRH | Neurot | -0.080 | 0.023 | -0.115 | -0.044 | 4.45e-04 | -0.014 | 0.034 | -0.081 | 0.052 | 0.671 | -0.023 | 82.200 | 0.167 |
| SRH | SCZ | -0.043 | 0.023 | -0.078 | -0.008 | 0.057 | -0.038 | 0.035 | -0.108 | 0.031 | 0.267 | -0.039 | 10.710 | 0.682 |
| SRH | SRH | 0.138 | 0.022 | 0.104 | 0.172 | 1.06e-09 | 0.076 | 0.035 | 0.004 | 0.148 | 0.03 | 0.084 | 45.160 | 0.323 |

**Note.** BMI = Body Mass Index; IQ = Intelligence; GCSE = General Certificate of Secondary Education (educational achievement); ADHD = Attention-Deficit/Hyperactivity Disorder; SCZ = Schizophrenia symptoms; EA = Educational Attainment; Neurot = Neuroticism; SRH = Self-rated Health; B = Between-family estimate; W = Within-family estimate; P = p-value of estimate; TotEff = Total effect derived as the intra-class correlation weighted sum of the within- and between family effect. PercReduc = Reduction of prediction estimates when comparing within- to between-family estimates in percentage. P.diff = empirical significance of difference between within- and between-family estimates based on permutation testing with 100,000 iterations.

Table S5. Within- and between-family prediction estimates after accounting for family socio-economic status

| Phenotype | GPS | beta.B | SE.B | L.CI.B | U.CI.B | P.B | beta.W | SE.W | CI.L.W | CI.U.W | P.W | TotEff | PercReduc | P.diff |
|---|---|---|---|---|---|---|---|---|---|---|---|---|---|---|
| ADHD | ADHD | 0.087 | 0.019 | 0.061 | 0.114 | 4.54e-06 | 0.122 | 0.024 | 0.072 | 0.174 | 2.50e-07 | 0.111 | 39.790 | 0.206 |
| ADHD | BMI | 0.057 | 0.019 | 0.032 | 0.083 | 0.002 | 0.041 | 0.024 | -0.010 | 0.093 | 0.088 | 0.046 | 28.760 | 0.52 |
| ADHD | EA | -0.051 | 0.020 | -0.079 | -0.024 | 0.01 | -0.139 | 0.025 | -0.196 | -0.082 | 4.62e-08 | -0.110 | 171.340 | 6.20e-04 |
| ADHD | Height | -0.013 | 0.018 | -0.040 | 0.013 | 0.488 | 0.021 | 0.024 | -0.030 | 0.072 | 0.374 | 0.010 | 265.070 | 0.32 |
| ADHD | IQ | -0.067 | 0.019 | -0.093 | -0.042 | 3.79e-04 | -0.112 | 0.024 | -0.163 | -0.062 | 4.63e-06 | -0.098 | 67.120 | 0.079 |
| ADHD | Neurot | 0.031 | 0.019 | 0.006 | 0.058 | 0.096 | 0.055 | 0.023 | 0.005 | 0.104 | 0.017 | 0.047 | 74.950 | 0.457 |
| ADHD | SCZ | 0.002 | 0.019 | -0.023 | 0.028 | 0.922 | 0.038 | 0.024 | -0.012 | 0.091 | 0.112 | 0.026 | 2,006.880 | 0.155 |
| ADHD | SRH | -0.087 | 0.019 | -0.113 | -0.060 | 5.60e-06 | -0.033 | 0.025 | -0.086 | 0.019 | 0.183 | -0.050 | 62.050 | 0.034 |
| BMI | ADHD | 0.041 | 0.025 | 0.006 | 0.077 | 0.107 | 0.054 | 0.032 | -0.015 | 0.123 | 0.093 | 0.050 | 32.390 | 0.785 |
| BMI | BMI | 0.328 | 0.023 | 0.294 | 0.362 | 1.10e-41 | 0.299 | 0.031 | 0.234 | 0.364 | 2.79e-21 | 0.308 | 9.010 | 0.226 |
| BMI | EA | -0.039 | 0.027 | -0.078 | -0.001 | 0.15 | -0.063 | 0.035 | -0.138 | 0.013 | 0.07 | -0.055 | 60.150 | 0.703 |
| BMI | Height | -0.046 | 0.024 | -0.080 | -0.013 | 0.059 | -0.030 | 0.032 | -0.102 | 0.042 | 0.35 | -0.035 | 34.520 | 0.798 |
| BMI | IQ | -0.010 | 0.025 | -0.044 | 0.025 | 0.703 | 0.014 | 0.033 | -0.061 | 0.088 | 0.671 | 0.007 | 245.160 | 0.583 |
| BMI | Neurot | -0.085 | 0.025 | -0.120 | -0.049 | 8.14e-04 | 0.004 | 0.032 | -0.062 | 0.069 | 0.91 | -0.025 | 104.250 | 0.034 |
| BMI | SCZ | -0.049 | 0.025 | -0.083 | -0.014 | 0.053 | -0.053 | 0.033 | -0.122 | 0.015 | 0.102 | -0.052 | 9.350 | 0.463 |
| BMI | SRH | -0.102 | 0.026 | -0.140 | -0.065 | 9.73e-05 | -0.142 | 0.032 | -0.207 | -0.074 | 1.31e-05 | -0.129 | 38.440 | 0.203 |
| GCSE | ADHD | -0.083 | 0.018 | -0.105 | -0.061 | 5.40e-06 | -0.062 | 0.019 | -0.104 | -0.018 | 0.001 | -0.074 | 25.620 | 0.406 |
| GCSE | BMI | -0.056 | 0.018 | -0.079 | -0.034 | 0.002 | -0.024 | 0.019 | -0.068 | 0.020 | 0.226 | -0.042 | 57.860 | 0.139 |
| GCSE | EA | 0.243 | 0.019 | 0.220 | 0.265 | 5.39e-38 | 0.210 | 0.020 | 0.165 | 0.257 | 3.36e-25 | 0.229 | 13.430 | 0.241 |
| GCSE | Height | 0.008 | 0.018 | -0.015 | 0.031 | 0.639 | -0.002 | 0.019 | -0.047 | 0.044 | 0.914 | 0.004 | 125.270 | 0.602 |
| GCSE | IQ | 0.212 | 0.018 | 0.191 | 0.233 | 8.86e-32 | 0.199 | 0.019 | 0.155 | 0.244 | 5.02e-24 | 0.207 | 5.980 | 0.461 |
| GCSE | Neurot | -0.037 | 0.018 | -0.059 | -0.015 | 0.043 | -0.063 | 0.019 | -0.105 | -0.020 | 7.27e-04 | -0.048 | 69.830 | 0.299 |
| GCSE | SCZ | 0.029 | 0.018 | 0.007 | 0.051 | 0.101 | -0.018 | 0.019 | -0.063 | 0.028 | 0.354 | 0.009 | 161.650 | 0.051 |

| | | | | | | | | | | | | | |
|---|---|---|---|---|---|---|---|---|---|---|---|---|---|
| GCSE | SRH | 0.069 | 0.019 | 0.046 | 0.093 | 1.87e-04 | 0.055 | 0.020 | 0.010 | 0.100 | 0.005 | 0.063 | 21.280 | 0.462 |
| Height | ADHD | -0.026 | 0.027 | -0.060 | 0.008 | 0.332 | -0.061 | 0.028 | -0.125 | 0.004 | 0.031 | -0.045 | 136.370 | 0.104 |
| Height | BMI | -0.017 | 0.026 | -0.050 | 0.016 | 0.523 | 0.013 | 0.028 | -0.050 | 0.074 | 0.637 | 0 | 179.180 | 0.582 |
| Height | EA | 0.013 | 0.028 | -0.023 | 0.049 | 0.645 | -0.002 | 0.030 | -0.072 | 0.068 | 0.96 | 0.005 | 111.660 | 0.479 |
| Height | Height | 0.474 | 0.022 | 0.446 | 0.500 | 2.94e-91 | 0.399 | 0.026 | 0.339 | 0.461 | 1.56e-49 | 0.432 | 15.630 | 0.017 |
| Height | IQ | 0.027 | 0.026 | -0.007 | 0.060 | 0.307 | 0.003 | 0.029 | -0.061 | 0.067 | 0.906 | 0.014 | 87.230 | 0.309 |
| Height | Neurot | -0.055 | 0.026 | -0.089 | -0.021 | 0.035 | 0.009 | 0.028 | -0.055 | 0.076 | 0.734 | -0.019 | 117.140 | 0.141 |
| Height | SCZ | -0.053 | 0.026 | -0.087 | -0.020 | 0.042 | 0.012 | 0.028 | -0.052 | 0.077 | 0.67 | -0.016 | 122.950 | 0.128 |
| Height | SRH | 0.047 | 0.027 | 0.014 | 0.080 | 0.082 | -0.020 | 0.029 | -0.081 | 0.042 | 0.493 | 0.010 | 141.700 | 0.339 |
| IQ | ADHD | -0.041 | 0.023 | -0.071 | -0.009 | 0.077 | -0.003 | 0.028 | -0.065 | 0.059 | 0.923 | -0.019 | 93.340 | 0.303 |
| IQ | BMI | 0.022 | 0.023 | -0.009 | 0.051 | 0.346 | 0.009 | 0.028 | -0.052 | 0.071 | 0.751 | 0.014 | 59.530 | 0.436 |
| IQ | EA | 0.128 | 0.024 | 0.096 | 0.160 | 1.28e-07 | 0.119 | 0.030 | 0.051 | 0.187 | 8.18e-05 | 0.122 | 7.050 | 0.786 |
| IQ | Height | -0.006 | 0.023 | -0.036 | 0.025 | 0.81 | 0.041 | 0.027 | -0.022 | 0.106 | 0.128 | 0.022 | 850.220 | 0.258 |
| IQ | IQ | 0.191 | 0.023 | 0.161 | 0.222 | 3.28e-16 | 0.129 | 0.029 | 0.067 | 0.192 | 6.67e-06 | 0.155 | 32.520 | 0.129 |
| IQ | Neurot | 0.002 | 0.023 | -0.029 | 0.033 | 0.928 | -0.030 | 0.027 | -0.090 | 0.033 | 0.269 | -0.016 | 1,526.480 | 0.455 |
| IQ | SCZ | 0.008 | 0.023 | -0.022 | 0.038 | 0.735 | 0.012 | 0.028 | -0.049 | 0.073 | 0.673 | 0.010 | 48.290 | 0.407 |
| IQ | SRH | 0.032 | 0.023 | 0.001 | 0.064 | 0.175 | 0.025 | 0.029 | -0.039 | 0.088 | 0.391 | 0.028 | 22.040 | 0.986 |
| Neurot | ADHD | -0.013 | 0.032 | -0.064 | 0.037 | 0.686 | 0.083 | 0.048 | -0.007 | 0.174 | 0.087 | 0.073 | 742.070 | 0.27 |
| Neurot | BMI | -0.020 | 0.032 | -0.072 | 0.032 | 0.531 | 0.018 | 0.047 | -0.082 | 0.114 | 0.708 | 0.014 | 189.260 | 0.383 |
| Neurot | EA | 0.030 | 0.032 | -0.022 | 0.082 | 0.337 | 0.024 | 0.055 | -0.088 | 0.138 | 0.654 | 0.025 | 19.320 | 0.851 |
| Neurot | Height | -0.021 | 0.031 | -0.072 | 0.029 | 0.487 | 0.035 | 0.048 | -0.058 | 0.124 | 0.469 | 0.029 | 262.920 | 0.227 |
| Neurot | IQ | -0.002 | 0.032 | -0.051 | 0.048 | 0.959 | 0.074 | 0.050 | -0.027 | 0.176 | 0.138 | 0.066 | 4,652.280 | 0.11 |
| Neurot | Neurot | 0.109 | 0.032 | 0.056 | 0.163 | 6.90e-04 | 0.013 | 0.047 | -0.083 | 0.114 | 0.779 | 0.023 | 87.790 | 0.039 |
| Neurot | SCZ | -0.012 | 0.031 | -0.060 | 0.035 | 0.689 | 0.053 | 0.049 | -0.046 | 0.153 | 0.284 | 0.046 | 522.340 | 0.255 |
| Neurot | SRH | -0.034 | 0.031 | -0.083 | 0.015 | 0.283 | -0.014 | 0.050 | -0.119 | 0.086 | 0.775 | -0.016 | 58.020 | 0.527 |

| | | | | | | | | | | | | | | |
|---|---|---|---|---|---|---|---|---|---|---|---|---|---|---|
| SCZ | ADHD | 0.029 | 0.028 | -0.008 | 0.068 | 0.297 | 0.005 | 0.037 | -0.076 | 0.083 | 0.892 | 0.011 | 82.660 | 0.914 |
| SCZ | BMI | 0.101 | 0.027 | 0.062 | 0.140 | 1.80e-04 | 0.043 | 0.037 | -0.037 | 0.125 | 0.25 | 0.057 | 57.440 | 0.235 |
| SCZ | EA | -0.049 | 0.029 | -0.093 | -0.005 | 0.09 | 0.014 | 0.040 | -0.071 | 0.099 | 0.731 | -0.002 | 127.820 | 0.093 |
| SCZ | Height | -0.006 | 0.026 | -0.044 | 0.032 | 0.81 | -0.002 | 0.037 | -0.074 | 0.069 | 0.958 | -0.003 | 68.920 | 0.878 |
| SCZ | IQ | 0.002 | 0.027 | -0.038 | 0.041 | 0.953 | 0.021 | 0.039 | -0.058 | 0.102 | 0.582 | 0.016 | 1,242.260 | 0.865 |
| SCZ | Neurot | 0.055 | 0.028 | 0.018 | 0.092 | 0.047 | -0.037 | 0.036 | -0.105 | 0.033 | 0.305 | -0.013 | 166.680 | 0.051 |
| SCZ | SCZ | -0.004 | 0.027 | -0.044 | 0.036 | 0.878 | -0.017 | 0.038 | -0.098 | 0.064 | 0.653 | -0.014 | 306.850 | 0.954 |
| SCZ | SRH | -0.100 | 0.028 | -0.139 | -0.062 | 3.28e-04 | -0.005 | 0.037 | -0.082 | 0.071 | 0.892 | -0.029 | 94.960 | 0.053 |
| SRH | ADHD | -0.046 | 0.023 | -0.082 | -0.011 | 0.046 | -0.074 | 0.035 | -0.145 | -0.004 | 0.032 | -0.071 | 60.590 | 0.805 |
| SRH | BMI | -0.105 | 0.023 | -0.140 | -0.070 | 4.04e-06 | -0.014 | 0.035 | -0.081 | 0.055 | 0.688 | -0.026 | 86.760 | 0.163 |
| SRH | EA | 0.052 | 0.024 | 0.015 | 0.090 | 0.033 | 0.078 | 0.037 | 0.005 | 0.153 | 0.037 | 0.074 | 48.780 | 0.392 |
| SRH | Height | 0.058 | 0.022 | 0.023 | 0.092 | 0.009 | -0.007 | 0.035 | -0.077 | 0.064 | 0.829 | 0.001 | 112.960 | 0.097 |
| SRH | IQ | 0.007 | 0.023 | -0.029 | 0.043 | 0.762 | 0.030 | 0.036 | -0.046 | 0.106 | 0.396 | 0.027 | 336.400 | 0.118 |
| SRH | Neurot | -0.073 | 0.023 | -0.109 | -0.036 | 0.002 | -0.016 | 0.034 | -0.083 | 0.051 | 0.632 | -0.024 | 77.470 | 0.26 |
| SRH | SCZ | -0.041 | 0.023 | -0.076 | -0.005 | 0.071 | -0.040 | 0.035 | -0.112 | 0.032 | 0.261 | -0.040 | 3.180 | 0.669 |
| SRH | SRH | 0.128 | 0.023 | 0.092 | 0.164 | 4.15e-08 | 0.075 | 0.035 | 0.003 | 0.148 | 0.034 | 0.082 | 41.880 | 0.486 |

**Note.** BMI = Body Mass Index; IQ = Intelligence; GCSE = General Certificate of Secondary Education (educational achievement); ADHD = Attention-Deficit/Hyperactivity Disorder; SCZ = Schizophrenia symptoms; EA = Educational Attainment; Neurot = Neuroticism; SRH = Self-rated Health; B = Between-family estimate; W = Within-family estimate; P = p-value of estimate; TotEff = Total effect derived as the intra-class correlation weighted sum of the within- and between family effect. PercReduc = Reduction of prediction estimates when comparing within- to between-family estimates in percentage. P.diff = empirical significance of difference between within- and between-family estimates based on permutation testing with 100,000 iterations.

Table S6. Within- and between-family prediction estimates for same-sex twin pairs

| Phenotype | GPS | beta.B | SE.B | L.CI.B | U.CI.B | P.B | beta.W | SE.W | CI.L.W | CI.U.W | P.W | TotEff | PercReduc | P.diff |
|---|---|---|---|---|---|---|---|---|---|---|---|---|---|---|
| ADHD | ADHD | 0.132 | 0.026 | 0.096 | 0.167 | 6.80e-07 | 0.165 | 0.033 | 0.092 | 0.239 | 5.86e-07 | 0.154 | 24.890 | 0.426 |
| ADHD | BMI | 0.088 | 0.026 | 0.053 | 0.124 | 6.93e-04 | 0.044 | 0.032 | -0.023 | 0.111 | 0.165 | 0.059 | 49.890 | 0.2 |
| ADHD | EA | -0.122 | 0.025 | -0.157 | -0.086 | 1.90e-06 | -0.209 | 0.034 | -0.286 | -0.132 | 9.68e-10 | -0.180 | 71.860 | 0.015 |
| ADHD | Height | -0.067 | 0.026 | -0.100 | -0.032 | 0.009 | -0.013 | 0.033 | -0.085 | 0.059 | 0.707 | -0.030 | 81.190 | 0.205 |
| ADHD | IQ | -0.107 | 0.026 | -0.142 | -0.071 | 3.74e-05 | -0.119 | 0.034 | -0.191 | -0.046 | 5.43e-04 | -0.115 | 11.290 | 0.766 |
| ADHD | Neurot | 0.043 | 0.026 | 0.006 | 0.079 | 0.095 | 0.090 | 0.032 | 0.023 | 0.160 | 0.005 | 0.075 | 109.100 | 0.245 |
| ADHD | SCZ | 0.013 | 0.026 | -0.023 | 0.048 | 0.62 | 0.059 | 0.033 | -0.011 | 0.130 | 0.074 | 0.044 | 357.450 | 0.174 |
| ADHD | SRH | -0.133 | 0.026 | -0.167 | -0.097 | 2.80e-07 | -0.073 | 0.034 | -0.147 | 0 | 0.033 | -0.093 | 45.140 | 0.067 |
| BMI | ADHD | 0.077 | 0.035 | 0.031 | 0.124 | 0.029 | 0.031 | 0.043 | -0.064 | 0.129 | 0.47 | 0.047 | 59.550 | 0.434 |
| BMI | BMI | 0.330 | 0.033 | 0.284 | 0.375 | 1.16e-22 | 0.309 | 0.040 | 0.221 | 0.397 | 2.53e-14 | 0.316 | 6.140 | 0.343 |
| BMI | EA | -0.120 | 0.035 | -0.167 | -0.075 | 6.78e-04 | -0.073 | 0.045 | -0.171 | 0.026 | 0.101 | -0.089 | 39.180 | 0.302 |
| BMI | Height | -0.093 | 0.034 | -0.138 | -0.046 | 0.006 | -0.073 | 0.044 | -0.171 | 0.028 | 0.096 | -0.080 | 21.270 | 0.817 |
| BMI | IQ | -0.065 | 0.035 | -0.112 | -0.017 | 0.061 | 0.012 | 0.044 | -0.084 | 0.106 | 0.794 | -0.014 | 117.690 | 0.292 |
| BMI | Neurot | -0.056 | 0.035 | -0.101 | -0.012 | 0.108 | -0.004 | 0.042 | -0.098 | 0.087 | 0.92 | -0.022 | 92.500 | 0.591 |
| BMI | SCZ | -0.017 | 0.035 | -0.066 | 0.033 | 0.626 | -0.079 | 0.043 | -0.173 | 0.013 | 0.064 | -0.058 | 363.290 | 0.634 |
| BMI | SRH | -0.172 | 0.035 | -0.218 | -0.125 | 8.52e-07 | -0.148 | 0.044 | -0.243 | -0.059 | 7.60e-04 | -0.156 | 14.100 | 0.629 |
| GCSE | ADHD | -0.188 | 0.029 | -0.221 | -0.155 | 1.24e-10 | -0.061 | 0.027 | -0.125 | 0.005 | 0.025 | -0.135 | 67.470 | 2.70e-04 |
| GCSE | BMI | -0.163 | 0.029 | -0.197 | -0.130 | 3.37e-08 | -0.023 | 0.026 | -0.084 | 0.038 | 0.387 | -0.104 | 86.160 | 7.00e-05 |
| GCSE | EA | 0.414 | 0.027 | 0.385 | 0.444 | 3.49e-49 | 0.222 | 0.027 | 0.158 | 0.287 | 6.33e-16 | 0.333 | 46.250 | 9.99e-06 |
| GCSE | Height | 0.061 | 0.029 | 0.029 | 0.093 | 0.032 | 0.029 | 0.027 | -0.037 | 0.095 | 0.283 | 0.048 | 52.170 | 0.473 |
| GCSE | IQ | 0.335 | 0.028 | 0.305 | 0.364 | 9.93e-32 | 0.204 | 0.027 | 0.139 | 0.267 | 1.96e-13 | 0.280 | 39.200 | 8.00e-05 |
| GCSE | Neurot | -0.085 | 0.029 | -0.118 | -0.053 | 0.003 | -0.112 | 0.026 | -0.173 | -0.051 | 1.31e-05 | -0.097 | 31.760 | 0.357 |
| GCSE | SCZ | -0.015 | 0.029 | -0.047 | 0.018 | 0.599 | -0.005 | 0.027 | -0.071 | 0.061 | 0.861 | -0.011 | 69.240 | 0.755 |

| | | | | | | | | | | | | | |
|---|---|---|---|---|---|---|---|---|---|---|---|---|---|
| GCSE | SRH | 0.228 | 0.028 | 0.197 | 0.258 | 2.04e-15 | 0.067 | 0.027 | 0.001 | 0.132 | 0.014 | 0.160 | 70.550 | 9.99e-06 |
| Height | ADHD | -0.025 | 0.035 | -0.070 | 0.021 | 0.474 | -0.067 | 0.038 | -0.153 | 0.021 | 0.081 | -0.049 | 163.440 | 0.101 |
| Height | BMI | -0.026 | 0.035 | -0.068 | 0.016 | 0.452 | -0.007 | 0.037 | -0.091 | 0.075 | 0.845 | -0.015 | 72.100 | 0.867 |
| Height | EA | 0.033 | 0.035 | -0.011 | 0.077 | 0.351 | -0.032 | 0.040 | -0.122 | 0.061 | 0.421 | -0.004 | 197.810 | 0.104 |
| Height | Height | 0.435 | 0.030 | 0.398 | 0.471 | 6.39e-43 | 0.391 | 0.037 | 0.307 | 0.474 | 4.77e-25 | 0.410 | 10.170 | 0.29 |
| Height | IQ | 0.063 | 0.035 | 0.020 | 0.106 | 0.069 | -0.048 | 0.039 | -0.131 | 0.034 | 0.221 | 0 | 176.300 | 0.011 |
| Height | Neurot | -0.064 | 0.035 | -0.109 | -0.017 | 0.064 | 0.024 | 0.037 | -0.067 | 0.119 | 0.52 | -0.014 | 137.540 | 0.024 |
| Height | SCZ | -0.059 | 0.035 | -0.102 | -0.016 | 0.093 | 0.039 | 0.038 | -0.050 | 0.128 | 0.305 | -0.003 | 166.690 | 0.077 |
| Height | SRH | 0.054 | 0.035 | 0.011 | 0.097 | 0.12 | -0.024 | 0.040 | -0.106 | 0.059 | 0.551 | 0.010 | 143.770 | 0.22 |
| IQ | ADHD | -0.079 | 0.033 | -0.119 | -0.038 | 0.018 | -0.030 | 0.037 | -0.113 | 0.054 | 0.423 | -0.052 | 62.350 | 0.284 |
| IQ | BMI | -0.010 | 0.034 | -0.049 | 0.030 | 0.766 | 0.017 | 0.036 | -0.068 | 0.100 | 0.641 | 0.005 | 268.950 | 0.539 |
| IQ | EA | 0.270 | 0.033 | 0.228 | 0.310 | 5.47e-16 | 0.142 | 0.039 | 0.051 | 0.231 | 2.77e-04 | 0.199 | 47.470 | 0.005 |
| IQ | Height | 0.060 | 0.034 | 0.018 | 0.103 | 0.073 | 0.049 | 0.037 | -0.041 | 0.139 | 0.184 | 0.054 | 18.480 | 0.519 |
| IQ | IQ | 0.266 | 0.034 | 0.225 | 0.306 | 1.25e-14 | 0.146 | 0.039 | 0.058 | 0.227 | 1.69e-04 | 0.200 | 45.140 | 0.009 |
| IQ | Neurot | -0.010 | 0.034 | -0.051 | 0.032 | 0.772 | -0.082 | 0.036 | -0.171 | 0.007 | 0.024 | -0.050 | 745.200 | 0.119 |
| IQ | SCZ | -0.017 | 0.034 | -0.058 | 0.025 | 0.625 | 0.005 | 0.037 | -0.077 | 0.086 | 0.902 | -0.005 | 127.100 | 0.595 |
| IQ | SRH | 0.114 | 0.033 | 0.072 | 0.155 | 6.14e-04 | 0.025 | 0.038 | -0.062 | 0.113 | 0.517 | 0.065 | 78.180 | 0.067 |
| Neurot | ADHD | -0.025 | 0.043 | -0.093 | 0.041 | 0.552 | 0.059 | 0.066 | -0.059 | 0.177 | 0.372 | 0.052 | 331.160 | 0.393 |
| Neurot | BMI | -0.029 | 0.042 | -0.100 | 0.041 | 0.492 | 0.054 | 0.063 | -0.075 | 0.187 | 0.386 | 0.047 | 286.230 | 0.16 |
| Neurot | EA | 0.020 | 0.039 | -0.043 | 0.084 | 0.613 | 0.099 | 0.073 | -0.060 | 0.254 | 0.177 | 0.092 | 399.840 | 0.222 |
| Neurot | Height | -0.012 | 0.041 | -0.080 | 0.057 | 0.767 | 0.018 | 0.066 | -0.106 | 0.142 | 0.783 | 0.016 | 247.020 | 0.631 |
| Neurot | IQ | -0.031 | 0.040 | -0.096 | 0.035 | 0.446 | 0.023 | 0.071 | -0.117 | 0.168 | 0.749 | 0.018 | 173.550 | 0.389 |
| Neurot | Neurot | 0.126 | 0.041 | 0.056 | 0.197 | 0.002 | 0.045 | 0.064 | -0.085 | 0.176 | 0.483 | 0.052 | 64.320 | 0.274 |
| Neurot | SCZ | -0.036 | 0.041 | -0.101 | 0.029 | 0.387 | 0.106 | 0.068 | -0.029 | 0.240 | 0.119 | 0.094 | 396.420 | 0.175 |
| Neurot | SRH | -0.056 | 0.040 | -0.118 | 0.008 | 0.161 | -0.011 | 0.068 | -0.146 | 0.126 | 0.875 | -0.015 | 80.760 | 0.361 |

| | | | | | | | | | | | | | | |
|---|---|---|---|---|---|---|---|---|---|---|---|---|---|---|
| SCZ | ADHD | 0.071 | 0.038 | 0.020 | 0.121 | 0.065 | 0.016 | 0.048 | -0.090 | 0.122 | 0.737 | 0.033 | 76.970 | 0.455 |
| SCZ | BMI | 0.118 | 0.038 | 0.065 | 0.172 | 0.002 | 0.082 | 0.048 | -0.017 | 0.183 | 0.085 | 0.093 | 30.160 | 0.276 |
| SCZ | EA | -0.064 | 0.037 | -0.117 | -0.014 | 0.083 | -0.015 | 0.049 | -0.122 | 0.094 | 0.763 | -0.030 | 76.860 | 0.199 |
| SCZ | Height | 0.022 | 0.036 | -0.027 | 0.070 | 0.553 | -0.020 | 0.049 | -0.118 | 0.079 | 0.685 | -0.007 | 192.280 | 0.629 |
| SCZ | IQ | -0.033 | 0.036 | -0.084 | 0.017 | 0.371 | 0.014 | 0.049 | -0.095 | 0.124 | 0.77 | 0 | 144.020 | 0.565 |
| SCZ | Neurot | 0.044 | 0.038 | -0.007 | 0.094 | 0.249 | 0.020 | 0.046 | -0.076 | 0.115 | 0.667 | 0.027 | 55.020 | 0.697 |
| SCZ | SCZ | 0.029 | 0.038 | -0.027 | 0.085 | 0.446 | 0.032 | 0.048 | -0.073 | 0.136 | 0.504 | 0.031 | 12.430 | 0.934 |
| SCZ | SRH | -0.134 | 0.038 | -0.186 | -0.084 | 4.49e-04 | -0.086 | 0.049 | -0.185 | 0.010 | 0.076 | -0.101 | 35.480 | 0.395 |
| SRH | ADHD | -0.068 | 0.032 | -0.117 | -0.018 | 0.033 | -0.086 | 0.046 | -0.179 | 0.004 | 0.059 | -0.083 | 27.450 | 0.735 |
| SRH | BMI | -0.132 | 0.031 | -0.177 | -0.086 | 2.23e-05 | -0.032 | 0.044 | -0.122 | 0.055 | 0.466 | -0.051 | 75.450 | 0.085 |
| SRH | EA | 0.065 | 0.031 | 0.018 | 0.111 | 0.04 | 0.073 | 0.047 | -0.023 | 0.168 | 0.121 | 0.072 | 13.650 | 0.96 |
| SRH | Height | 0.063 | 0.030 | 0.016 | 0.110 | 0.039 | 0.011 | 0.047 | -0.081 | 0.105 | 0.813 | 0.020 | 82.350 | 0.254 |
| SRH | IQ | 0.049 | 0.031 | 0.003 | 0.094 | 0.118 | 0.043 | 0.047 | -0.056 | 0.143 | 0.352 | 0.044 | 10.700 | 0.917 |
| SRH | Neurot | -0.089 | 0.031 | -0.136 | -0.042 | 0.004 | -0.025 | 0.044 | -0.117 | 0.069 | 0.568 | -0.037 | 71.450 | 0.123 |
| SRH | SCZ | -0.078 | 0.031 | -0.128 | -0.031 | 0.013 | -0.030 | 0.045 | -0.123 | 0.063 | 0.508 | -0.039 | 61.620 | 0.275 |
| SRH | SRH | 0.133 | 0.031 | 0.086 | 0.179 | 1.96e-05 | 0.129 | 0.047 | 0.028 | 0.232 | 0.006 | 0.130 | 2.930 | 0.94 |

**Note.** BMI = Body Mass Index; IQ = Intelligence; GCSE = General Certificate of Secondary Education (educational achievement); ADHD = Attention-Deficit/Hyperactivity Disorder; SCZ = Schizophrenia symptoms; EA = Educational Attainment; Neurot = Neuroticism; SRH = Self-rated Health; B = Between-family estimate; W = Within-family estimate; P = p-value of estimate; TotEff = Total effect derived as the intra-class correlation weighted sum of the within- and between family effect. PercReduc = Reduction of prediction estimates when comparing within- to between-family estimates in percentage. P.diff = empirical significance of difference between within- and between-family estimates based on permutation testing with 100,000 iterations.

Table S7. Within- and between-family prediction estimates after accounting for family socio-economic status for same-sex twin pairs

| Phenotype | GPS | beta.B | SE.B | L.CI.B | U.CI.B | P.B | beta.W | SE.W | CI.L.W | CI.U.W | P.W | TotEff | PercReduc | P.diff |
|---|---|---|---|---|---|---|---|---|---|---|---|---|---|---|
| ADHD | ADHD | 0.114 | 0.026 | 0.078 | 0.150 | 1.54e-05 | 0.168 | 0.033 | 0.094 | 0.242 | 5.04e-07 | 0.150 | 47.810 | 0.157 |
| ADHD | BMI | 0.057 | 0.026 | 0.020 | 0.093 | 0.029 | 0.042 | 0.032 | -0.025 | 0.109 | 0.197 | 0.047 | 25.770 | 0.639 |
| ADHD | EA | -0.065 | 0.027 | -0.102 | -0.027 | 0.018 | -0.208 | 0.034 | -0.287 | -0.130 | 2.07e-09 | -0.161 | 221.410 | 8.00e-05 |
| ADHD | Height | -0.049 | 0.026 | -0.083 | -0.015 | 0.056 | -0.006 | 0.034 | -0.080 | 0.067 | 0.861 | -0.020 | 87.890 | 0.357 |
| ADHD | IQ | -0.070 | 0.026 | -0.107 | -0.033 | 0.008 | -0.117 | 0.035 | -0.189 | -0.043 | 7.95e-04 | -0.102 | 66.610 | 0.189 |
| ADHD | Neurot | 0.038 | 0.025 | 0.001 | 0.075 | 0.136 | 0.096 | 0.033 | 0.025 | 0.165 | 0.004 | 0.077 | 151.340 | 0.135 |
| ADHD | SCZ | 0.014 | 0.026 | -0.022 | 0.050 | 0.576 | 0.055 | 0.033 | -0.019 | 0.126 | 0.098 | 0.042 | 288.780 | 0.225 |
| ADHD | SRH | -0.100 | 0.026 | -0.135 | -0.064 | 1.39e-04 | -0.069 | 0.035 | -0.140 | 0.004 | 0.048 | -0.079 | 31.150 | 0.314 |
| BMI | ADHD | 0.043 | 0.035 | -0.005 | 0.090 | 0.215 | 0.023 | 0.043 | -0.072 | 0.119 | 0.592 | 0.030 | 46.080 | 0.778 |
| BMI | BMI | 0.309 | 0.032 | 0.263 | 0.354 | 1.59e-20 | 0.313 | 0.040 | 0.225 | 0.398 | 1.44e-14 | 0.311 | 1.350 | 0.823 |
| BMI | EA | -0.050 | 0.037 | -0.100 | 0.001 | 0.179 | -0.063 | 0.045 | -0.161 | 0.036 | 0.161 | -0.058 | 24.560 | 0.788 |
| BMI | Height | -0.073 | 0.033 | -0.118 | -0.028 | 0.028 | -0.068 | 0.044 | -0.171 | 0.035 | 0.124 | -0.070 | 7.550 | 0.614 |
| BMI | IQ | -0.027 | 0.035 | -0.075 | 0.022 | 0.438 | 0.015 | 0.044 | -0.081 | 0.114 | 0.733 | 0.001 | 155.640 | 0.827 |
| BMI | Neurot | -0.067 | 0.034 | -0.111 | -0.022 | 0.049 | -0.005 | 0.043 | -0.099 | 0.086 | 0.907 | -0.026 | 92.560 | 0.413 |
| BMI | SCZ | -0.023 | 0.034 | -0.072 | 0.027 | 0.503 | -0.078 | 0.043 | -0.172 | 0.012 | 0.069 | -0.060 | 240.460 | 0.703 |
| BMI | SRH | -0.127 | 0.035 | -0.175 | -0.078 | 3.34e-04 | -0.156 | 0.044 | -0.252 | -0.064 | 3.72e-04 | -0.146 | 23.130 | 0.579 |
| GCSE | ADHD | -0.097 | 0.025 | -0.128 | -0.067 | 1.27e-04 | -0.072 | 0.028 | -0.135 | -0.010 | 0.01 | -0.086 | 25.660 | 0.524 |
| GCSE | BMI | -0.089 | 0.025 | -0.120 | -0.057 | 4.87e-04 | -0.021 | 0.027 | -0.079 | 0.039 | 0.425 | -0.060 | 76.080 | 0.039 |
| GCSE | EA | 0.247 | 0.026 | 0.216 | 0.278 | 7.23e-21 | 0.222 | 0.028 | 0.158 | 0.286 | 2.43e-15 | 0.236 | 10.320 | 0.486 |
| GCSE | Height | 0.015 | 0.025 | -0.015 | 0.046 | 0.539 | 0.030 | 0.028 | -0.035 | 0.094 | 0.288 | 0.021 | 94.320 | 0.599 |
| GCSE | IQ | 0.224 | 0.025 | 0.195 | 0.252 | 1.06e-18 | 0.207 | 0.028 | 0.142 | 0.272 | 2.36e-13 | 0.217 | 7.450 | 0.545 |
| GCSE | Neurot | -0.063 | 0.025 | -0.093 | -0.033 | 0.011 | -0.115 | 0.026 | -0.174 | -0.056 | 1.52e-05 | -0.085 | 82.830 | 0.075 |
| GCSE | SCZ | -0.005 | 0.025 | -0.036 | 0.025 | 0.84 | -0.002 | 0.027 | -0.067 | 0.062 | 0.927 | -0.004 | 50.010 | 0.926 |

| | | | | | | | | | | | | | |
|---|---|---|---|---|---|---|---|---|---|---|---|---|---|
| GCSE | SRH | 0.110 | 0.025 | 0.078 | 0.140 | 1.70e-05 | 0.067 | 0.028 | 0.002 | 0.131 | 0.017 | 0.091 | 39.200 | 0.251 |
| Height | ADHD | -0.013 | 0.036 | -0.058 | 0.034 | 0.717 | -0.067 | 0.039 | -0.155 | 0.022 | 0.084 | -0.044 | 418.390 | 0.044 |
| Height | BMI | -0.015 | 0.035 | -0.058 | 0.028 | 0.678 | -0.013 | 0.038 | -0.097 | 0.071 | 0.729 | -0.014 | 11.120 | 0.681 |
| Height | EA | -0.004 | 0.038 | -0.052 | 0.044 | 0.92 | -0.038 | 0.040 | -0.130 | 0.055 | 0.346 | -0.023 | 897.210 | 0.394 |
| Height | Height | 0.434 | 0.030 | 0.397 | 0.471 | 4.84e-42 | 0.386 | 0.037 | 0.301 | 0.470 | 7.41e-24 | 0.407 | 10.970 | 0.256 |
| Height | IQ | 0.040 | 0.036 | -0.005 | 0.083 | 0.266 | -0.050 | 0.040 | -0.135 | 0.034 | 0.212 | -0.011 | 224.500 | 0.029 |
| Height | Neurot | -0.064 | 0.035 | -0.111 | -0.017 | 0.066 | 0.033 | 0.038 | -0.057 | 0.128 | 0.382 | -0.009 | 152.190 | 0.039 |
| Height | SCZ | -0.056 | 0.035 | -0.100 | -0.010 | 0.11 | 0.036 | 0.039 | -0.056 | 0.127 | 0.359 | -0.004 | 163.310 | 0.085 |
| Height | SRH | 0.039 | 0.036 | -0.006 | 0.084 | 0.285 | -0.023 | 0.040 | -0.105 | 0.061 | 0.574 | 0.004 | 158.370 | 0.439 |
| IQ | ADHD | -0.040 | 0.031 | -0.080 | 0 | 0.202 | -0.031 | 0.038 | -0.115 | 0.053 | 0.414 | -0.035 | 22.330 | 0.983 |
| IQ | BMI | 0.025 | 0.032 | -0.014 | 0.064 | 0.433 | 0.019 | 0.037 | -0.066 | 0.105 | 0.607 | 0.022 | 22.720 | 0.726 |
| IQ | EA | 0.157 | 0.033 | 0.113 | 0.200 | 2.97e-06 | 0.132 | 0.039 | 0.043 | 0.222 | 8.53e-04 | 0.143 | 15.550 | 0.868 |
| IQ | Height | 0.019 | 0.032 | -0.022 | 0.059 | 0.56 | 0.041 | 0.038 | -0.047 | 0.131 | 0.273 | 0.031 | 121.640 | 0.758 |
| IQ | IQ | 0.199 | 0.033 | 0.158 | 0.241 | 2.21e-09 | 0.137 | 0.040 | 0.053 | 0.223 | 5.70e-04 | 0.165 | 31.390 | 0.258 |
| IQ | Neurot | 0.002 | 0.031 | -0.040 | 0.042 | 0.954 | -0.084 | 0.038 | -0.172 | 0.005 | 0.025 | -0.046 | 4,776 | 0.061 |
| IQ | SCZ | -0.006 | 0.032 | -0.046 | 0.035 | 0.853 | 0.008 | 0.038 | -0.076 | 0.091 | 0.836 | 0.002 | 231.510 | 0.617 |
| IQ | SRH | 0.046 | 0.032 | 0.004 | 0.088 | 0.146 | 0.021 | 0.039 | -0.067 | 0.110 | 0.594 | 0.032 | 55.100 | 0.864 |
| Neurot | ADHD | -0.029 | 0.044 | -0.101 | 0.039 | 0.499 | 0.063 | 0.067 | -0.056 | 0.182 | 0.348 | 0.055 | 312.800 | 0.297 |
| Neurot | BMI | -0.033 | 0.044 | -0.105 | 0.040 | 0.455 | 0.050 | 0.064 | -0.080 | 0.182 | 0.429 | 0.043 | 254.860 | 0.239 |
| Neurot | EA | 0.034 | 0.043 | -0.035 | 0.106 | 0.429 | 0.096 | 0.074 | -0.063 | 0.258 | 0.196 | 0.091 | 181.030 | 0.347 |
| Neurot | Height | -0.009 | 0.042 | -0.080 | 0.060 | 0.826 | 0.016 | 0.067 | -0.112 | 0.142 | 0.809 | 0.014 | 273.610 | 0.587 |
| Neurot | IQ | -0.030 | 0.043 | -0.101 | 0.038 | 0.481 | 0.030 | 0.072 | -0.113 | 0.173 | 0.678 | 0.025 | 198.500 | 0.354 |
| Neurot | Neurot | 0.130 | 0.042 | 0.060 | 0.201 | 0.002 | 0.041 | 0.066 | -0.095 | 0.175 | 0.536 | 0.048 | 68.540 | 0.196 |
| Neurot | SCZ | -0.041 | 0.042 | -0.106 | 0.024 | 0.334 | 0.090 | 0.069 | -0.045 | 0.225 | 0.194 | 0.079 | 321.750 | 0.177 |
| Neurot | SRH | -0.056 | 0.041 | -0.120 | 0.010 | 0.181 | -0.010 | 0.069 | -0.145 | 0.127 | 0.89 | -0.013 | 82.750 | 0.457 |

| | | | | | | | | | | | | | |
|---|---|---|---|---|---|---|---|---|---|---|---|---|---|
| SCZ | ADHD | 0.058 | 0.038 | 0.010 | 0.109 | 0.125 | 0.018 | 0.049 | -0.088 | 0.124 | 0.718 | 0.030 | 69.510 | 0.628 |
| SCZ | BMI | 0.103 | 0.038 | 0.050 | 0.158 | 0.006 | 0.091 | 0.048 | -0.009 | 0.192 | 0.06 | 0.095 | 12.020 | 0.504 |
| SCZ | EA | -0.044 | 0.039 | -0.100 | 0.013 | 0.268 | -0.018 | 0.050 | -0.128 | 0.096 | 0.72 | -0.026 | 58.820 | 0.436 |
| SCZ | Height | 0.034 | 0.036 | -0.016 | 0.086 | 0.343 | -0.021 | 0.050 | -0.122 | 0.080 | 0.667 | -0.004 | 162.630 | 0.559 |
| SCZ | IQ | -0.015 | 0.037 | -0.066 | 0.036 | 0.689 | 0.011 | 0.050 | -0.101 | 0.123 | 0.826 | 0.003 | 173.690 | 0.957 |
| SCZ | Neurot | 0.038 | 0.038 | -0.013 | 0.089 | 0.318 | 0.014 | 0.048 | -0.083 | 0.112 | 0.767 | 0.022 | 62.850 | 0.681 |
| SCZ | SCZ | 0.025 | 0.038 | -0.029 | 0.080 | 0.51 | 0.031 | 0.049 | -0.079 | 0.135 | 0.536 | 0.029 | 23.500 | 0.955 |
| SCZ | SRH | -0.113 | 0.039 | -0.166 | -0.060 | 0.003 | -0.084 | 0.049 | -0.180 | 0.012 | 0.088 | -0.093 | 25.870 | 0.634 |
| SRH | ADHD | -0.051 | 0.032 | -0.099 | -0.001 | 0.114 | -0.085 | 0.046 | -0.177 | 0.007 | 0.066 | -0.079 | 66.990 | 0.99 |
| SRH | BMI | -0.125 | 0.031 | -0.171 | -0.079 | 8.31e-05 | -0.036 | 0.045 | -0.126 | 0.053 | 0.426 | -0.052 | 71.250 | 0.168 |
| SRH | EA | 0.038 | 0.034 | -0.015 | 0.088 | 0.265 | 0.078 | 0.048 | -0.018 | 0.176 | 0.104 | 0.070 | 105.540 | 0.354 |
| SRH | Height | 0.052 | 0.030 | 0.007 | 0.099 | 0.086 | 0.014 | 0.047 | -0.078 | 0.105 | 0.76 | 0.021 | 72.330 | 0.384 |
| SRH | IQ | 0.028 | 0.032 | -0.019 | 0.076 | 0.38 | 0.050 | 0.047 | -0.050 | 0.149 | 0.287 | 0.046 | 77.880 | 0.414 |
| SRH | Neurot | -0.086 | 0.031 | -0.134 | -0.037 | 0.006 | -0.027 | 0.045 | -0.123 | 0.067 | 0.55 | -0.038 | 68.280 | 0.232 |
| SRH | SCZ | -0.067 | 0.032 | -0.117 | -0.018 | 0.033 | -0.036 | 0.046 | -0.128 | 0.059 | 0.439 | -0.042 | 46.960 | 0.486 |
| SRH | SRH | 0.122 | 0.032 | 0.073 | 0.170 | 1.53e-04 | 0.131 | 0.048 | 0.028 | 0.236 | 0.006 | 0.129 | 6.970 | 0.906 |

**Note.** BMI = Body Mass Index; IQ = Intelligence; GCSE = General Certificate of Secondary Education (educational achievement); ADHD = Attention-Deficit/Hyperactivity Disorder; SCZ = Schizophrenia symptoms; EA = Educational Attainment; Neurot = Neuroticism; SRH = Self-rated Health; B = Between-family estimate; W = Within-family estimate; P = p-value of estimate; TotEff = Total effect derived as the intra-class correlation weighted sum of the within- and between family effect. PercReduc = Reduction of prediction estimates when comparing within- to between-family estimates in percentage. P.diff = empirical significance of difference between within- and between-family estimates based on permutation testing with 100,000 iterations.

Table S8. Intraclass coefficients and residual effects for same-sex and opposite-sex twin pairs

| | Same-Sex Twin Pairs | | | | | Opposite-Sex Twin Pairs | | | | |
|---|---|---|---|---|---|---|---|---|---|---|
| **Phenotype** | **N pairs** | **ICC** | **ICC 95% CI lower** | **ICC 95% CI upper** | **RandEff resid** | **N pairs** | **ICC** | **ICC 95% CI lower** | **ICC 95% CI upper** | **RandEff resid** |
| **Height** | 789 | 0.435 | 0.365 | 0.52 | 0.565 | 674 | 0.442 | 0.367 | 0.533 | 0.557 |
| **BMI** | 733 | 0.339 | 0.271 | 0.424 | 0.661 | 620 | 0.286 | 0.215 | 0.381 | 0.713 |
| **Self-rated Health** | 805 | 0.182 | 0.123 | 0.268 | 0.818 | 689 | 0.083 | 0.031 | 0.221 | 0.916 |
| **IQ** | 824 | 0.450 | 0.381 | 0.531 | 0.549 | 745 | 0.386 | 0.313 | 0.471 | 0.614 |
| **GCSE** | 1,220 | 0.579 | 0.518 | 0.647 | 0.421 | 1,146 | 0.585 | 0.521 | 0.655 | 0.415 |
| **Neuroticism** | 429 | 0.084 | 0.027 | 0.262 | 0.915 | 360 | 0.123 | 0.052 | 0.288 | 0.876 |
| **ADHD Symptoms** | 1,285 | 0.328 | 0.278 | 0.390 | 0.671 | 1,184 | 0.317 | 0.263 | 0.381 | 0.683 |
| **SCZ Symptoms** | 613 | 0.308 | 0.235 | 0.402 | 0.692 | 527 | 0.194 | 0.124 | 0.301 | 0.805 |

**Note.** BMI = Body Mass Index; IQ = Intelligence; GCSE = General Certificate of Secondary Education (educational achievement); ADHD = Attention-Deficit/Hyperactivity Disorder; SCZ = Schizophrenia symptoms; EA = Educational Attainment; Neurot = Neuroticism; ICC = Intraclass coefficient; CI = Confidence Interval; RandEff Resid = Residual of the random effect.

Table S9. Within- and between-family prediction estimates for opposite-sex twin pairs

| Phenotype | GPS | beta.B | SE.B | L.CI.B | U.CI.B | P.B | beta.W | SE.W | CI.L.W | CI.U.W | P.W | TotEff | PercReduc | P.diff |
|---|---|---|---|---|---|---|---|---|---|---|---|---|---|---|
| ADHD | ADHD | 0.087 | 0.028 | 0.050 | 0.124 | 0.002 | 0.081 | 0.034 | 0.008 | 0.154 | 0.017 | 0.083 | 7.650 | 0.88 |
| ADHD | BMI | 0.099 | 0.027 | 0.060 | 0.137 | 2.64e-04 | 0.029 | 0.036 | -0.047 | 0.104 | 0.415 | 0.051 | 70.610 | 0.079 |
| ADHD | EA | -0.114 | 0.026 | -0.149 | -0.078 | 1.77e-05 | -0.057 | 0.038 | -0.141 | 0.028 | 0.132 | -0.075 | 50.360 | 0.163 |
| ADHD | Height | 0.035 | 0.027 | -0.004 | 0.074 | 0.197 | 0.049 | 0.034 | -0.022 | 0.119 | 0.15 | 0.044 | 38.190 | 0.833 |
| ADHD | IQ | -0.105 | 0.027 | -0.142 | -0.070 | 8.45e-05 | -0.125 | 0.035 | -0.198 | -0.051 | 3.36e-04 | -0.119 | 18.310 | 0.611 |
| ADHD | Neurot | 0.046 | 0.029 | 0.009 | 0.082 | 0.109 | 0.013 | 0.033 | -0.058 | 0.082 | 0.687 | 0.024 | 71.170 | 0.344 |
| ADHD | SCZ | -0.017 | 0.028 | -0.053 | 0.018 | 0.533 | 0.015 | 0.035 | -0.059 | 0.089 | 0.659 | 0.005 | 189.840 | 0.433 |
| ADHD | SRH | -0.116 | 0.027 | -0.153 | -0.078 | 2.15e-05 | 0.005 | 0.035 | -0.073 | 0.084 | 0.882 | -0.033 | 104.500 | 0.002 |
| BMI | ADHD | 0.055 | 0.038 | 0.004 | 0.106 | 0.148 | 0.094 | 0.048 | -0.004 | 0.194 | 0.049 | 0.083 | 72.020 | 0.792 |
| BMI | BMI | 0.375 | 0.034 | 0.325 | 0.425 | 7.22e-26 | 0.279 | 0.048 | 0.184 | 0.374 | 1.12e-08 | 0.306 | 25.660 | 0.061 |
| BMI | EA | -0.103 | 0.036 | -0.158 | -0.048 | 0.005 | -0.057 | 0.054 | -0.171 | 0.060 | 0.293 | -0.070 | 45.250 | 0.291 |
| BMI | Height | 0.011 | 0.036 | -0.039 | 0.062 | 0.764 | 0.014 | 0.047 | -0.086 | 0.116 | 0.757 | 0.013 | 32.910 | 0.486 |
| BMI | IQ | -0.027 | 0.036 | -0.075 | 0.021 | 0.455 | 0.008 | 0.049 | -0.107 | 0.124 | 0.869 | -0.002 | 129.740 | 0.343 |
| BMI | Neurot | -0.113 | 0.038 | -0.169 | -0.058 | 0.003 | -0.011 | 0.047 | -0.104 | 0.083 | 0.821 | -0.040 | 90.630 | 0.101 |
| BMI | SCZ | -0.092 | 0.038 | -0.140 | -0.044 | 0.015 | -0.024 | 0.049 | -0.125 | 0.078 | 0.618 | -0.044 | 73.520 | 0.182 |
| BMI | SRH | -0.102 | 0.038 | -0.160 | -0.046 | 0.008 | -0.117 | 0.048 | -0.214 | -0.023 | 0.014 | -0.113 | 14.890 | 0.632 |
| GCSE | ADHD | -0.164 | 0.030 | -0.198 | -0.130 | 8.52e-08 | -0.061 | 0.027 | -0.122 | -0.001 | 0.024 | -0.121 | 62.870 | 0.003 |
| GCSE | BMI | -0.106 | 0.030 | -0.139 | -0.073 | 4.84e-04 | -0.023 | 0.028 | -0.089 | 0.044 | 0.424 | -0.072 | 78.620 | 0.009 |
| GCSE | EA | 0.423 | 0.027 | 0.392 | 0.453 | 4.30e-51 | 0.204 | 0.029 | 0.137 | 0.271 | 4.48e-12 | 0.332 | 51.780 | 1.00e-05 |
| GCSE | Height | 0.003 | 0.030 | -0.033 | 0.039 | 0.91 | -0.029 | 0.027 | -0.095 | 0.035 | 0.277 | -0.010 | 973.950 | 0.233 |
| GCSE | IQ | 0.305 | 0.029 | 0.273 | 0.336 | 4.22e-25 | 0.198 | 0.027 | 0.137 | 0.261 | 5.86e-13 | 0.261 | 35 | 7.70e-04 |
| GCSE | Neurot | -0.062 | 0.032 | -0.097 | -0.027 | 0.052 | -0.009 | 0.026 | -0.072 | 0.051 | 0.719 | -0.040 | 84.940 | 0.066 |
| GCSE | SCZ | 0.098 | 0.030 | 0.066 | 0.131 | 0.001 | -0.030 | 0.027 | -0.096 | 0.038 | 0.278 | 0.045 | 130.250 | 3.10e-04 |

| | | | | | | | | | | | | | | |
|---|---|---|---|---|---|---|---|---|---|---|---|---|---|---|
| GCSE | SRH | 0.144 | 0.031 | 0.107 | 0.180 | 3.98e-06 | 0.048 | 0.028 | -0.019 | 0.113 | 0.084 | 0.104 | 66.870 | 0.002 |
| Height | ADHD | -0.047 | 0.039 | -0.095 | 0.002 | 0.228 | -0.046 | 0.041 | -0.142 | 0.051 | 0.263 | -0.046 | 1.820 | 0.967 |
| Height | BMI | -0.015 | 0.038 | -0.065 | 0.035 | 0.689 | 0.064 | 0.042 | -0.026 | 0.153 | 0.124 | 0.029 | 521.910 | 0.112 |
| Height | EA | 0.036 | 0.037 | -0.011 | 0.082 | 0.338 | 0.032 | 0.046 | -0.074 | 0.135 | 0.493 | 0.033 | 11.830 | 0.792 |
| Height | Height | 0.500 | 0.031 | 0.458 | 0.545 | 3.83e-49 | 0.429 | 0.036 | 0.341 | 0.517 | 1.35e-29 | 0.461 | 14.260 | 0.159 |
| Height | IQ | 0.019 | 0.037 | -0.029 | 0.067 | 0.609 | 0.039 | 0.042 | -0.056 | 0.135 | 0.346 | 0.030 | 110.250 | 0.697 |
| Height | Neurot | -0.051 | 0.039 | -0.099 | -0.004 | 0.19 | -0.012 | 0.040 | -0.099 | 0.076 | 0.764 | -0.029 | 76.380 | 0.859 |
| Height | SCZ | -0.043 | 0.038 | -0.092 | 0.006 | 0.257 | -0.012 | 0.041 | -0.099 | 0.075 | 0.779 | -0.025 | 72.920 | 0.611 |
| Height | SRH | 0.061 | 0.039 | 0.016 | 0.107 | 0.115 | -0.030 | 0.041 | -0.117 | 0.058 | 0.461 | 0.010 | 149.070 | 0.403 |
| IQ | ADHD | -0.095 | 0.036 | -0.142 | -0.046 | 0.008 | 0.024 | 0.041 | -0.063 | 0.114 | 0.549 | -0.022 | 125.780 | 0.014 |
| IQ | BMI | -0.052 | 0.035 | -0.099 | -0.006 | 0.143 | 0.011 | 0.041 | -0.078 | 0.103 | 0.787 | -0.013 | 121.300 | 0.322 |
| IQ | EA | 0.236 | 0.033 | 0.191 | 0.280 | 2.89e-12 | 0.106 | 0.046 | 0.005 | 0.207 | 0.021 | 0.156 | 54.800 | 0.003 |
| IQ | Height | -0.052 | 0.036 | -0.098 | -0.007 | 0.144 | 0.040 | 0.039 | -0.053 | 0.131 | 0.317 | 0.004 | 175.930 | 0.135 |
| IQ | IQ | 0.259 | 0.033 | 0.217 | 0.303 | 3.11e-14 | 0.127 | 0.041 | 0.040 | 0.216 | 0.002 | 0.178 | 50.920 | 0.015 |
| IQ | Neurot | -0.032 | 0.036 | -0.082 | 0.018 | 0.381 | 0.048 | 0.038 | -0.035 | 0.132 | 0.208 | 0.017 | 251.490 | 0.052 |
| IQ | SCZ | 0.033 | 0.036 | -0.013 | 0.079 | 0.365 | 0.021 | 0.041 | -0.070 | 0.111 | 0.603 | 0.026 | 35.040 | 0.539 |
| IQ | SRH | 0.104 | 0.036 | 0.056 | 0.150 | 0.004 | 0.019 | 0.043 | -0.075 | 0.112 | 0.651 | 0.052 | 81.280 | 0.055 |
| Neurot | ADHD | 0.005 | 0.046 | -0.064 | 0.075 | 0.909 | 0.106 | 0.070 | -0.030 | 0.245 | 0.131 | 0.094 | 1,915.360 | 0.618 |
| Neurot | BMI | -0.004 | 0.046 | -0.076 | 0.068 | 0.923 | -0.023 | 0.071 | -0.173 | 0.123 | 0.742 | -0.021 | 428.580 | 0.975 |
| Neurot | EA | 0.011 | 0.043 | -0.059 | 0.082 | 0.789 | -0.082 | 0.079 | -0.234 | 0.073 | 0.294 | -0.071 | 824.020 | 0.149 |
| Neurot | Height | -0.037 | 0.044 | -0.105 | 0.031 | 0.393 | 0.055 | 0.068 | -0.084 | 0.193 | 0.427 | 0.043 | 246.220 | 0.257 |
| Neurot | IQ | 0.026 | 0.046 | -0.044 | 0.096 | 0.572 | 0.090 | 0.069 | -0.058 | 0.239 | 0.193 | 0.082 | 247.440 | 0.414 |
| Neurot | Neurot | 0.087 | 0.049 | 0.005 | 0.171 | 0.076 | -0.006 | 0.068 | -0.148 | 0.134 | 0.932 | 0.006 | 106.600 | 0.175 |
| Neurot | SCZ | 0.013 | 0.046 | -0.055 | 0.083 | 0.782 | 0.018 | 0.070 | -0.135 | 0.169 | 0.791 | 0.018 | 43.330 | 0.697 |
| Neurot | SRH | -0.020 | 0.046 | -0.090 | 0.052 | 0.667 | -0.039 | 0.070 | -0.190 | 0.110 | 0.579 | -0.036 | 96.100 | 0.999 |

| | | | | | | | | | | | | | |
|---|---|---|---|---|---|---|---|---|---|---|---|---|---|
| SCZ | ADHD | 0.012 | 0.041 | -0.045 | 0.069 | 0.773 | -0.012 | 0.055 | -0.128 | 0.106 | 0.823 | -0.008 | 204.090 | 0.883 |
| SCZ | BMI | 0.099 | 0.038 | 0.044 | 0.155 | 0.01 | -0.010 | 0.057 | -0.136 | 0.123 | 0.861 | 0.011 | 110.120 | 0.323 |
| SCZ | EA | -0.102 | 0.039 | -0.163 | -0.042 | 0.01 | 0.043 | 0.063 | -0.088 | 0.171 | 0.499 | 0.015 | 142.060 | 0.029 |
| SCZ | Height | -0.026 | 0.038 | -0.088 | 0.036 | 0.496 | 0.026 | 0.055 | -0.081 | 0.130 | 0.64 | 0.016 | 201.050 | 0.583 |
| SCZ | IQ | -0.013 | 0.039 | -0.073 | 0.046 | 0.731 | 0.039 | 0.060 | -0.077 | 0.158 | 0.52 | 0.029 | 389.350 | 0.432 |
| SCZ | Neurot | 0.082 | 0.040 | 0.024 | 0.138 | 0.043 | -0.070 | 0.053 | -0.170 | 0.033 | 0.182 | -0.041 | 185.910 | 0.04 |
| SCZ | SCZ | -0.058 | 0.040 | -0.112 | -0.003 | 0.148 | -0.060 | 0.057 | -0.183 | 0.063 | 0.289 | -0.060 | 4.980 | 0.721 |
| SCZ | SRH | -0.110 | 0.040 | -0.165 | -0.056 | 0.006 | 0.069 | 0.055 | -0.050 | 0.189 | 0.214 | 0.034 | 162.530 | 0.011 |
| SRH | ADHD | -0.042 | 0.033 | -0.093 | 0.010 | 0.206 | -0.054 | 0.052 | -0.160 | 0.052 | 0.298 | -0.053 | 29.160 | 0.861 |
| SRH | BMI | -0.076 | 0.032 | -0.129 | -0.025 | 0.019 | 0.016 | 0.053 | -0.092 | 0.123 | 0.756 | 0.009 | 121.540 | 0.602 |
| SRH | EA | 0.075 | 0.032 | 0.024 | 0.124 | 0.02 | 0.087 | 0.058 | -0.025 | 0.199 | 0.134 | 0.086 | 16.590 | 0.892 |
| SRH | Height | 0.061 | 0.031 | 0.011 | 0.112 | 0.052 | -0.029 | 0.050 | -0.130 | 0.074 | 0.569 | -0.021 | 146.860 | 0.111 |
| SRH | IQ | -0.007 | 0.032 | -0.060 | 0.046 | 0.822 | 0.008 | 0.053 | -0.105 | 0.118 | 0.886 | 0.006 | 206.960 | 0.218 |
| SRH | Neurot | -0.069 | 0.034 | -0.122 | -0.014 | 0.041 | -0.001 | 0.051 | -0.098 | 0.094 | 0.98 | -0.007 | 98.180 | 0.669 |
| SRH | SCZ | -0.003 | 0.032 | -0.054 | 0.048 | 0.922 | -0.048 | 0.053 | -0.153 | 0.059 | 0.362 | -0.044 | 1,423.010 | 0.098 |
| SRH | SRH | 0.144 | 0.033 | 0.093 | 0.194 | 1.32e-05 | 0.020 | 0.052 | -0.082 | 0.119 | 0.705 | 0.030 | 86.400 | 0.236 |

**Note.** BMI = Body Mass Index; IQ = Intelligence; GCSE = General Certificate of Secondary Education (educational achievement); ADHD = Attention-Deficit/Hyperactivity Disorder; SCZ = Schizophrenia symptoms; EA = Educational Attainment; Neurot = Neuroticism; SRH = Self-rated Health; B = Between-family estimate; W = Within-family estimate; P = p-value of estimate; TotEff = Total effect derived as the intra-class correlation weighted sum of the within- and between family effect. PercReduc = Reduction of prediction estimates when comparing within- to between-family estimates in percentage. P.diff = empirical significance of difference between within- and between-family estimates based on permutation testing with 100,000 iterations.

Table S10. Within- and between-family prediction estimates after accounting for family socio-economic status for opposite-sex twin pairs

| Phenotype | GPS | beta.B | SE.B | L.CI.B | U.CI.B | P.B | beta.W | SE.W | CI.L.W | CI.U.W | P.W | TotEff | PercReduc | P.diff |
|---|---|---|---|---|---|---|---|---|---|---|---|---|---|---|
| ADHD | ADHD | 0.056 | 0.028 | 0.018 | 0.095 | 0.042 | 0.074 | 0.033 | 0.0004 | 0.144 | 0.028 | 0.068 | 30.950 | 0.639 |
| ADHD | BMI | 0.059 | 0.027 | 0.021 | 0.096 | 0.03 | 0.039 | 0.035 | -0.037 | 0.115 | 0.267 | 0.045 | 32.970 | 0.660 |
| ADHD | EA | -0.034 | 0.029 | -0.075 | 0.007 | 0.241 | -0.055 | 0.037 | -0.140 | 0.029 | 0.14 | -0.048 | 61.930 | 0.492 |
| ADHD | Height | 0.027 | 0.027 | -0.014 | 0.067 | 0.319 | 0.050 | 0.034 | -0.021 | 0.122 | 0.139 | 0.042 | 86.080 | 0.662 |
| ADHD | IQ | -0.064 | 0.027 | -0.102 | -0.025 | 0.017 | -0.108 | 0.034 | -0.177 | -0.036 | 0.002 | -0.094 | 66.990 | 0.244 |
| ADHD | Neurot | 0.022 | 0.028 | -0.014 | 0.059 | 0.429 | 0.014 | 0.032 | -0.056 | 0.084 | 0.662 | 0.017 | 36.680 | 0.749 |
| ADHD | SCZ | -0.013 | 0.027 | -0.047 | 0.023 | 0.64 | 0.019 | 0.035 | -0.056 | 0.092 | 0.593 | 0.009 | 246.760 | 0.427 |
| ADHD | SRH | -0.071 | 0.028 | -0.109 | -0.033 | 0.011 | 0.006 | 0.035 | -0.073 | 0.083 | 0.865 | -0.018 | 108.360 | 0.052 |
| BMI | ADHD | 0.038 | 0.038 | -0.015 | 0.092 | 0.312 | 0.093 | 0.049 | -0.008 | 0.193 | 0.056 | 0.077 | 144.380 | 0.518 |
| BMI | BMI | 0.354 | 0.034 | 0.303 | 0.403 | 2.49e-23 | 0.277 | 0.049 | 0.182 | 0.375 | 2.77e-08 | 0.299 | 21.660 | 0.119 |
| BMI | EA | -0.027 | 0.040 | -0.087 | 0.037 | 0.507 | -0.063 | 0.055 | -0.180 | 0.054 | 0.25 | -0.052 | 137.050 | 0.743 |
| BMI | Height | -0.010 | 0.036 | -0.062 | 0.040 | 0.773 | 0.012 | 0.047 | -0.090 | 0.115 | 0.8 | 0.006 | 215.540 | 0.456 |
| BMI | IQ | 0.011 | 0.037 | -0.040 | 0.062 | 0.758 | 0.013 | 0.050 | -0.105 | 0.127 | 0.798 | 0.012 | 13.050 | 0.595 |
| BMI | Neurot | -0.110 | 0.038 | -0.165 | -0.054 | 0.004 | 0.014 | 0.048 | -0.076 | 0.107 | 0.766 | -0.021 | 113.120 | 0.032 |
| BMI | SCZ | -0.083 | 0.037 | -0.133 | -0.034 | 0.026 | -0.021 | 0.050 | -0.124 | 0.082 | 0.683 | -0.038 | 75.260 | 0.171 |
| BMI | SRH | -0.069 | 0.039 | -0.128 | -0.010 | 0.08 | -0.125 | 0.048 | -0.222 | -0.029 | 0.01 | -0.109 | 81.810 | 0.202 |
| GCSE | ADHD | -0.067 | 0.026 | -0.100 | -0.035 | 0.011 | -0.052 | 0.027 | -0.108 | 0.006 | 0.053 | -0.061 | 22.960 | 0.642 |
| GCSE | BMI | -0.023 | 0.026 | -0.054 | 0.007 | 0.38 | -0.026 | 0.029 | -0.094 | 0.040 | 0.356 | -0.024 | 15.650 | 0.980 |
| GCSE | EA | 0.238 | 0.027 | 0.205 | 0.270 | 1.41e-18 | 0.198 | 0.029 | 0.132 | 0.261 | 2.01e-11 | 0.221 | 16.860 | 0.330 |
| GCSE | Height | 0.002 | 0.025 | -0.032 | 0.035 | 0.947 | -0.033 | 0.027 | -0.095 | 0.029 | 0.222 | -0.013 | 2,047.970 | 0.205 |
| GCSE | IQ | 0.199 | 0.025 | 0.168 | 0.230 | 9.70e-15 | 0.191 | 0.027 | 0.132 | 0.252 | 3.32e-12 | 0.196 | 4.070 | 0.676 |
| GCSE | Neurot | -0.007 | 0.027 | -0.039 | 0.025 | 0.8 | -0.012 | 0.026 | -0.072 | 0.049 | 0.656 | -0.009 | 68.750 | 0.883 |

| | | | | | | | | | | | | | | |
|---|---|---|---|---|---|---|---|---|---|---|---|---|---|---|
| GCSE | SCZ | 0.066 | 0.026 | 0.035 | 0.097 | 0.011 | -0.034 | 0.028 | -0.098 | 0.030 | 0.214 | 0.024 | 152.310 | 0.003 |
| GCSE | SRH | 0.025 | 0.027 | -0.010 | 0.059 | 0.36 | 0.043 | 0.028 | -0.021 | 0.106 | 0.122 | 0.032 | 71.170 | 0.824 |
| Height | ADHD | -0.040 | 0.039 | -0.090 | 0.011 | 0.311 | -0.054 | 0.041 | -0.149 | 0.043 | 0.189 | -0.048 | 34.380 | 0.798 |
| Height | BMI | -0.017 | 0.039 | -0.066 | 0.034 | 0.658 | 0.046 | 0.042 | -0.042 | 0.137 | 0.277 | 0.018 | 365.720 | 0.200 |
| Height | EA | 0.035 | 0.042 | -0.017 | 0.087 | 0.4 | 0.046 | 0.046 | -0.063 | 0.150 | 0.32 | 0.041 | 30.510 | 0.920 |
| Height | Height | 0.519 | 0.031 | 0.478 | 0.559 | 2.98e-51 | 0.411 | 0.036 | 0.327 | 0.498 | 2.47e-27 | 0.459 | 20.670 | 0.026 |
| Height | IQ | 0.012 | 0.038 | -0.040 | 0.063 | 0.756 | 0.062 | 0.042 | -0.036 | 0.159 | 0.141 | 0.040 | 423.670 | 0.397 |
| Height | Neurot | -0.044 | 0.040 | -0.093 | 0.004 | 0.267 | -0.017 | 0.041 | -0.109 | 0.072 | 0.669 | -0.029 | 60.570 | 0.960 |
| Height | SCZ | -0.047 | 0.038 | -0.098 | 0.003 | 0.222 | -0.015 | 0.042 | -0.105 | 0.074 | 0.721 | -0.029 | 68.120 | 0.716 |
| Height | SRH | 0.055 | 0.040 | 0.006 | 0.103 | 0.173 | -0.016 | 0.041 | -0.106 | 0.072 | 0.686 | 0.015 | 129.830 | 0.674 |
| IQ | ADHD | -0.043 | 0.034 | -0.092 | 0.005 | 0.206 | 0.029 | 0.041 | -0.060 | 0.119 | 0.477 | 0.001 | 168.250 | 0.148 |
| IQ | BMI | 0.017 | 0.033 | -0.029 | 0.062 | 0.617 | -0.004 | 0.041 | -0.096 | 0.088 | 0.93 | 0.004 | 121.980 | 0.457 |
| IQ | EA | 0.094 | 0.035 | 0.047 | 0.144 | 0.007 | 0.101 | 0.046 | -0.001 | 0.202 | 0.03 | 0.098 | 7.500 | 0.791 |
| IQ | Height | -0.034 | 0.033 | -0.079 | 0.011 | 0.305 | 0.042 | 0.040 | -0.051 | 0.131 | 0.293 | 0.013 | 223.100 | 0.188 |
| IQ | IQ | 0.183 | 0.033 | 0.138 | 0.227 | 2.81e-08 | 0.122 | 0.042 | 0.033 | 0.209 | 0.004 | 0.145 | 33.500 | 0.331 |
| IQ | Neurot | -0.001 | 0.034 | -0.051 | 0.048 | 0.981 | 0.026 | 0.039 | -0.059 | 0.109 | 0.507 | 0.016 | 3,246.290 | 0.485 |
| IQ | SCZ | 0.025 | 0.034 | -0.021 | 0.069 | 0.449 | 0.016 | 0.041 | -0.075 | 0.105 | 0.695 | 0.020 | 36.990 | 0.516 |
| IQ | SRH | 0.015 | 0.034 | -0.033 | 0.063 | 0.656 | 0.029 | 0.043 | -0.064 | 0.120 | 0.494 | 0.024 | 91.590 | 0.853 |
| Neurot | ADHD | 0.002 | 0.047 | -0.069 | 0.075 | 0.97 | 0.106 | 0.070 | -0.034 | 0.242 | 0.134 | 0.093 | 5,910.410 | 0.617 |
| Neurot | BMI | -0.007 | 0.047 | -0.080 | 0.067 | 0.882 | -0.024 | 0.072 | -0.173 | 0.126 | 0.736 | -0.022 | 248.420 | 0.992 |
| Neurot | EA | 0.027 | 0.046 | -0.050 | 0.105 | 0.555 | -0.061 | 0.081 | -0.217 | 0.101 | 0.446 | -0.051 | 325.450 | 0.188 |
| Neurot | Height | -0.033 | 0.045 | -0.105 | 0.040 | 0.465 | 0.055 | 0.069 | -0.083 | 0.196 | 0.427 | 0.044 | 267.330 | 0.262 |
| Neurot | IQ | 0.033 | 0.047 | -0.037 | 0.102 | 0.486 | 0.118 | 0.070 | -0.030 | 0.268 | 0.093 | 0.107 | 259.050 | 0.235 |
| Neurot | Neurot | 0.085 | 0.050 | 0.002 | 0.169 | 0.091 | -0.017 | 0.068 | -0.160 | 0.126 | 0.807 | -0.004 | 119.700 | 0.145 |
| Neurot | SCZ | 0.020 | 0.047 | -0.052 | 0.090 | 0.668 | 0.014 | 0.070 | -0.139 | 0.163 | 0.842 | 0.015 | 30.540 | 0.793 |

| | | | | | | | | | | | | | | |
|---|---|---|---|---|---|---|---|---|---|---|---|---|---|---|
| Neurot | SRH | -0.004 | 0.048 | -0.078 | 0.071 | 0.939 | -0.019 | 0.071 | -0.179 | 0.135 | 0.789 | -0.017 | 415.930 | 0.943 |
| SCZ | ADHD | -0.006 | 0.041 | -0.065 | 0.055 | 0.891 | -0.009 | 0.056 | -0.125 | 0.112 | 0.873 | -0.008 | 58.700 | 0.517 |
| SCZ | BMI | 0.099 | 0.038 | 0.043 | 0.155 | 0.009 | -0.015 | 0.058 | -0.148 | 0.118 | 0.793 | 0.007 | 115.370 | 0.306 |
| SCZ | EA | -0.055 | 0.043 | -0.126 | 0.015 | 0.203 | 0.058 | 0.064 | -0.071 | 0.187 | 0.366 | 0.036 | 205.500 | 0.107 |
| SCZ | Height | -0.055 | 0.037 | -0.115 | 0.005 | 0.14 | 0.019 | 0.056 | -0.083 | 0.123 | 0.734 | 0.005 | 134.400 | 0.429 |
| SCZ | IQ | 0.022 | 0.039 | -0.042 | 0.085 | 0.572 | 0.034 | 0.060 | -0.085 | 0.153 | 0.573 | 0.032 | 52.990 | 0.844 |
| SCZ | Neurot | 0.074 | 0.040 | 0.019 | 0.129 | 0.064 | -0.090 | 0.053 | -0.187 | 0.011 | 0.091 | -0.058 | 221.340 | 0.026 |
| SCZ | SCZ | -0.040 | 0.039 | -0.096 | 0.015 | 0.313 | -0.073 | 0.058 | -0.198 | 0.050 | 0.208 | -0.067 | 84.020 | 0.965 |
| SCZ | SRH | -0.088 | 0.040 | -0.143 | -0.031 | 0.03 | 0.082 | 0.056 | -0.035 | 0.200 | 0.143 | 0.049 | 193.450 | 0.027 |
| SRH | ADHD | -0.040 | 0.034 | -0.092 | 0.012 | 0.232 | -0.062 | 0.053 | -0.172 | 0.047 | 0.237 | -0.060 | 55.490 | 0.876 |
| SRH | BMI | -0.081 | 0.033 | -0.133 | -0.026 | 0.014 | 0.014 | 0.054 | -0.094 | 0.122 | 0.794 | 0.006 | 117.420 | 0.342 |
| SRH | EA | 0.072 | 0.035 | 0.014 | 0.129 | 0.043 | 0.077 | 0.059 | -0.037 | 0.192 | 0.189 | 0.077 | 7.840 | 0.532 |
| SRH | Height | 0.061 | 0.032 | 0.009 | 0.113 | 0.057 | -0.030 | 0.051 | -0.133 | 0.072 | 0.553 | -0.023 | 149.830 | 0.161 |
| SRH | IQ | -0.017 | 0.032 | -0.071 | 0.036 | 0.6 | 0.007 | 0.054 | -0.110 | 0.121 | 0.898 | 0.005 | 140.750 | 0.150 |
| SRH | Neurot | -0.056 | 0.034 | -0.110 | 0.001 | 0.1 | -0.004 | 0.052 | -0.101 | 0.093 | 0.942 | -0.008 | 93.180 | 0.704 |
| SRH | SCZ | -0.010 | 0.032 | -0.062 | 0.043 | 0.76 | -0.044 | 0.054 | -0.155 | 0.068 | 0.413 | -0.041 | 345.920 | 0.197 |
| SRH | SRH | 0.135 | 0.034 | 0.080 | 0.187 | 7.66e-05 | 0.015 | 0.052 | -0.086 | 0.115 | 0.771 | 0.025 | 88.720 | 0.127 |

**Note.** BMI = Body Mass Index; IQ = Intelligence; GCSE = General Certificate of Secondary Education (educational achievement); ADHD = Attention-Deficit/Hyperactivity Disorder; SCZ = Schizophrenia symptoms; EA = Educational Attainment; Neurot = Neuroticism; SRH = Self-rated Health; B = Between-family estimate; W = Within-family estimate; P = p-value of estimate; TotEff = Total effect derived as the intra-class correlation weighted sum of the within- and between family effect. PercReduc = Reduction of prediction estimates when comparing within- to between-family estimates in percentage. P.diff = empirical significance of difference between within- and between-family estimates based on permutation testing with 100,000 iterations.

Table S11 Phenotypic and polygenic score mean differences by polygenic score difference quantiles

| phenotype | GPS quant | mean phen | CI.L | CI.U | mean GPS | CI.L | CI.U |
|---|---|---|---|---|---|---|---|
| Height | 1 | 0.270 | -1.339 | 1.880 | -0.004 | -0.016 | 0.008 |
| Height | 2 | 1.718 | 0.203 | 3.233 | -0.003 | -0.033 | 0.027 |
| Height | 3 | 2.435 | 0.738 | 4.132 | 0.020 | -0.030 | 0.071 |
| Height | 4 | 3.489 | 2.039 | 4.939 | -0.030 | -0.102 | 0.041 |
| Height | 5 | 1.433 | -0.209 | 3.075 | 0.066 | -0.025 | 0.158 |
| Height | 6 | 1.452 | -0.144 | 3.048 | 0.019 | -0.101 | 0.139 |
| Height | 7 | 4.620 | 2.862 | 6.379 | -0.007 | -0.154 | 0.141 |
| Height | 8 | 3.649 | 1.813 | 5.485 | -0.038 | -0.218 | 0.142 |
| Height | 9 | 7.021 | 5.316 | 8.727 | 0.065 | -0.165 | 0.295 |
| Height | 10 | 8.989 | 7.181 | 10.798 | -0.090 | -0.422 | 0.242 |
| BMI | 1 | 0.064 | -0.843 | 0.972 | 0.001 | -0.012 | 0.014 |
| BMI | 2 | 0.968 | 0.126 | 1.810 | 0.004 | -0.029 | 0.037 |
| BMI | 3 | -0.555 | -1.439 | 0.329 | -0.011 | -0.066 | 0.044 |
| BMI | 4 | 1.204 | 0.319 | 2.089 | -0.027 | -0.105 | 0.051 |
| BMI | 5 | 0.689 | -0.073 | 1.451 | 0.055 | -0.047 | 0.158 |
| BMI | 6 | 0.885 | -0.009 | 1.778 | -0.031 | -0.159 | 0.097 |
| BMI | 7 | 1.495 | 0.584 | 2.406 | 0.075 | -0.084 | 0.235 |
| BMI | 8 | 1.309 | 0.410 | 2.208 | -0.084 | -0.281 | 0.113 |
| BMI | 9 | 1.762 | 0.999 | 2.526 | 0.043 | -0.200 | 0.286 |
| BMI | 10 | 2.933 | 2.092 | 3.773 | -0.026 | -0.378 | 0.327 |
| IQ | 1 | 1.369 | -1.040 | 3.778 | 0.005 | -0.006 | 0.016 |
| IQ | 2 | -0.260 | -2.860 | 2.341 | -0.011 | -0.042 | 0.019 |
| IQ | 3 | 1.139 | -1.385 | 3.662 | 0.010 | -0.041 | 0.061 |
| IQ | 4 | -0.725 | -3.287 | 1.838 | -0.009 | -0.081 | 0.063 |
| IQ | 5 | 2.081 | -0.367 | 4.529 | 0.023 | -0.069 | 0.115 |
| IQ | 6 | 1.483 | -0.982 | 3.949 | 0.019 | -0.099 | 0.137 |
| IQ | 7 | 1.288 | -1.222 | 3.799 | -0.051 | -0.193 | 0.092 |
| IQ | 8 | 2.379 | -0.215 | 4.973 | 0.092 | -0.083 | 0.268 |
| IQ | 9 | 4.304 | 1.881 | 6.728 | -0.101 | -0.324 | 0.121 |
| IQ | 10 | 3.291 | 0.934 | 5.647 | 0.023 | -0.293 | 0.340 |
| GCSE | 1 | 0.068 | -0.067 | 0.203 | 0.005 | -0.004 | 0.013 |
| GCSE | 2 | -0.057 | -0.204 | 0.090 | 0.001 | -0.023 | 0.024 |
| GCSE | 3 | 0.195 | 0.053 | 0.336 | -0.012 | -0.051 | 0.026 |
| GCSE | 4 | 0.080 | -0.063 | 0.223 | 0.021 | -0.033 | 0.074 |
| GCSE | 5 | 0.097 | -0.039 | 0.234 | -0.022 | -0.094 | 0.049 |
| GCSE | 6 | 0.266 | 0.129 | 0.403 | 0.054 | -0.039 | 0.146 |

| | | | | | | | |
|---|---|---|---|---|---|---|---|
| GCSE | 7 | 0.226 | 0.083 | 0.369 | 0.084 | -0.030 | 0.199 |
| GCSE | 8 | 0.307 | 0.180 | 0.435 | 0.060 | -0.079 | 0.200 |
| GCSE | 9 | 0.341 | 0.192 | 0.490 | 0.013 | -0.159 | 0.185 |
| GCSE | 10 | 0.471 | 0.327 | 0.614 | -0.202 | -0.445 | 0.041 |

**Note.** BMI = Body Mass Index; IQ = Intelligence; GCSE = General Certificate of Secondary Education (educational achievement); GPS = genome wide polygenic score; quant = quantile; phen = phenotype; CI.L = 95% lower confidence interval; CI.U = 95% upper confidence interval; GPS quant 1 = lowest absolute GPS twin pair difference quantile; GPS quant 10 = highest absolute GPS twin pair difference quantile.

#### Supplementary Figures

Figure S1. Within-twin pair Pearson's correlation coefficients.

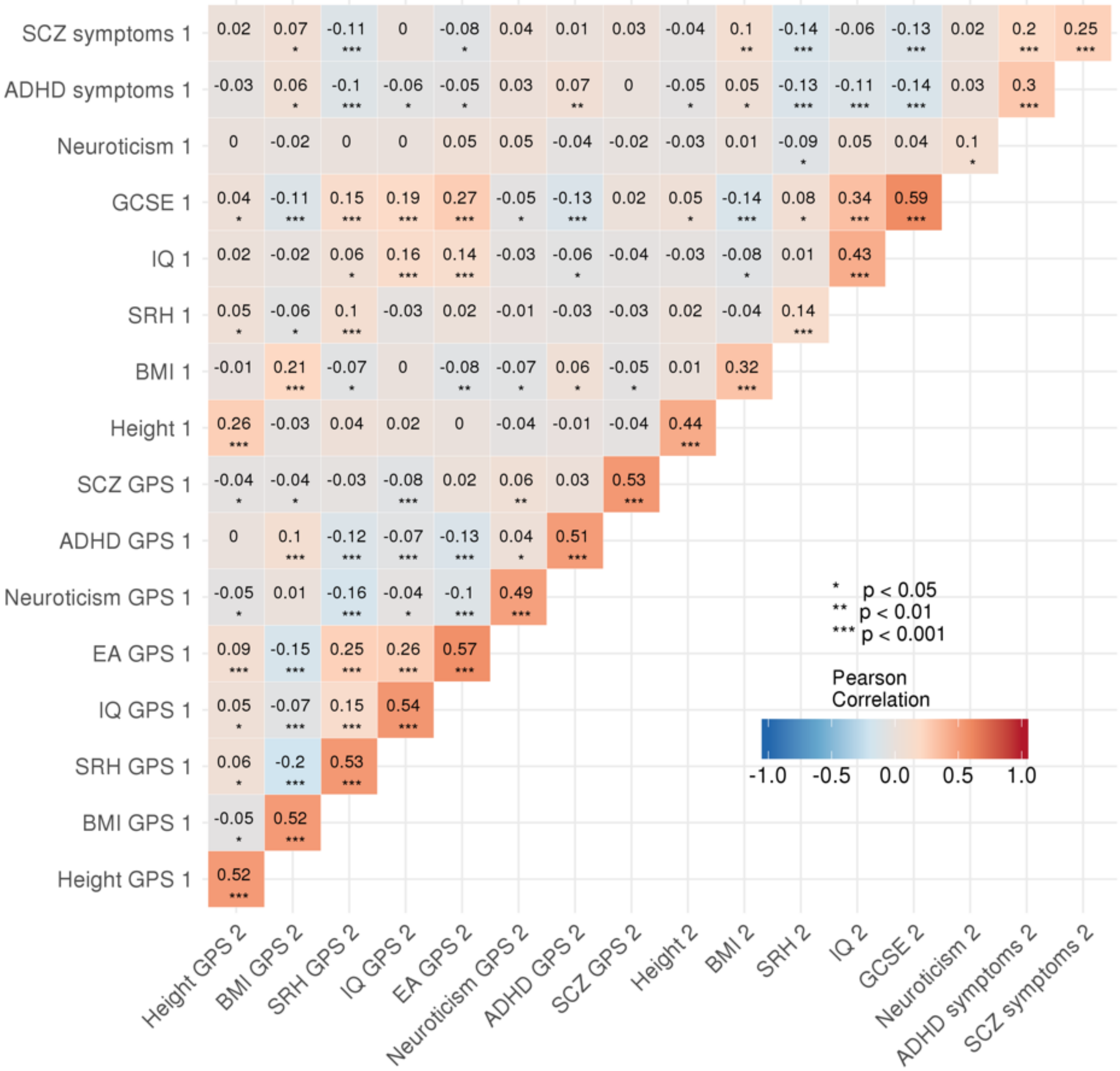

**Note.** BMI = Body Mass Index; IQ = Intelligence; GCSE = General Certificate of Secondary Education (educational achievement); ADHD = Attention-Deficit/Hyperactivity Disorder; SCZ = Schizophrenia; EA = Educational Attainment; SRH = Self-rated Health; 1 = Twin 1; 2 = Twin 2.

Figure S2. Within- and between-family prediction estimates of eight outcomes using eight polygenic scores for same-sex twin pairs

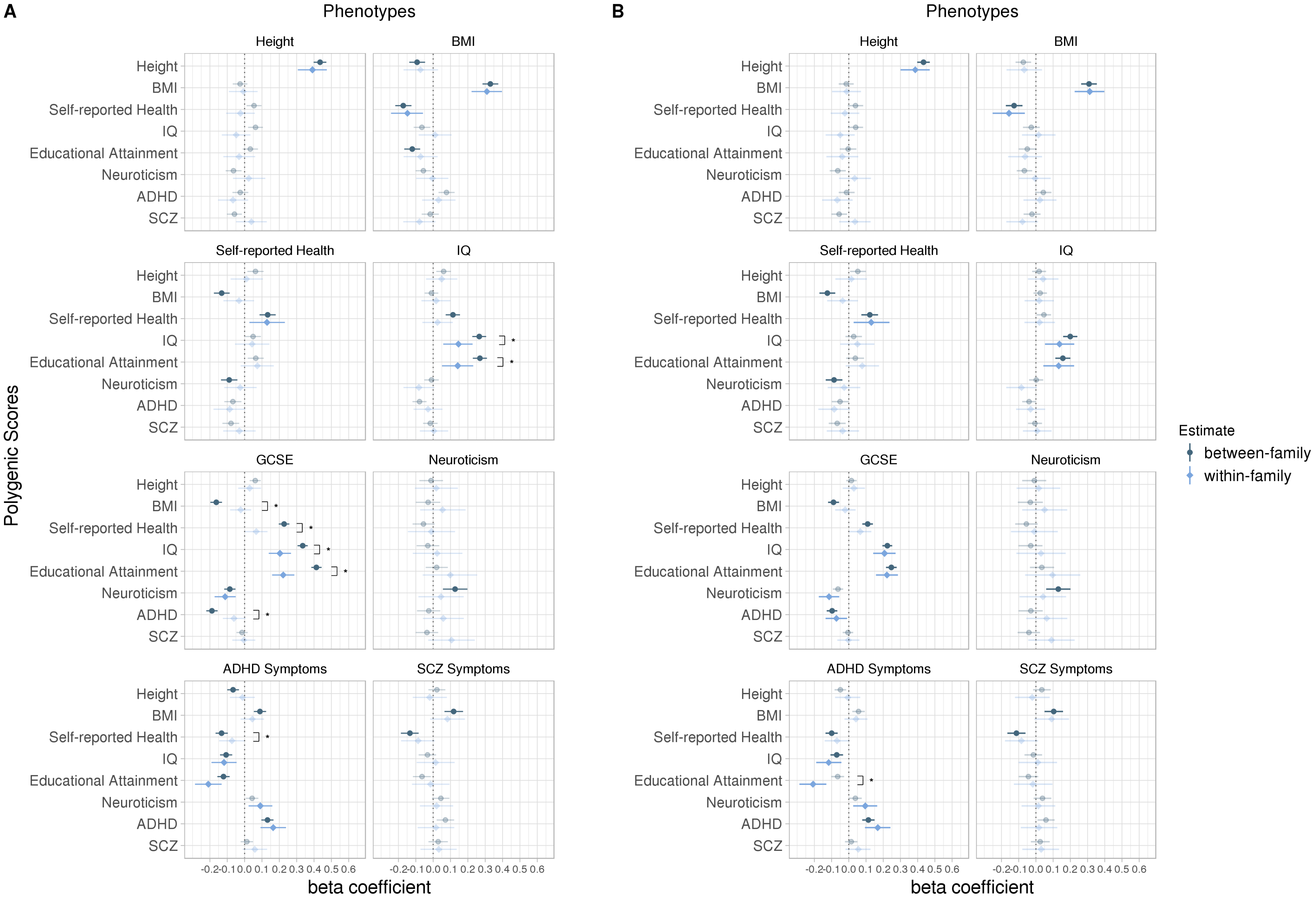

**Note.** Within- and between-family prediction estimates of eight developmental outcomes in same-sex twin pairs using eight polygenic scores before (A) and after (B) statistical correction for family socio-economic status, based on same-sex twin pairs only. Genome-wide Polygenic Scores are presented on the y-axis, predicting each of the eight phenotypic traits. Error bars are 95% bootstrap percentile intervals based on 10,000 bootstrap samples. Opaque estimates indicate statistical significance at the false discovery rate corrected threshold of $p < 0.01$. Brackets indicate a significant difference between within- and between-family prediction estimate based on permutation testing with 100,000 iterations, and only significant differences are shown where at least one of the estimates is significant at the false discovery rate corrected threshold of $p < 0.01$ (for all prediction estimates and statistical significance, see Supplementary Tables S6 and S7). The dotted line represents a beta coefficient of zero. BMI = Body Mass Index; IQ = Intelligence; GCSE = General Certificate of Secondary Education (educational achievement); ADHD = Attention-Deficit/Hyperactivity Disorder; SCZ = Schizophrenia.

Figure S3. Within- and between-family prediction estimates of eight outcomes using eight polygenic scores for opposite-sex twin pairs

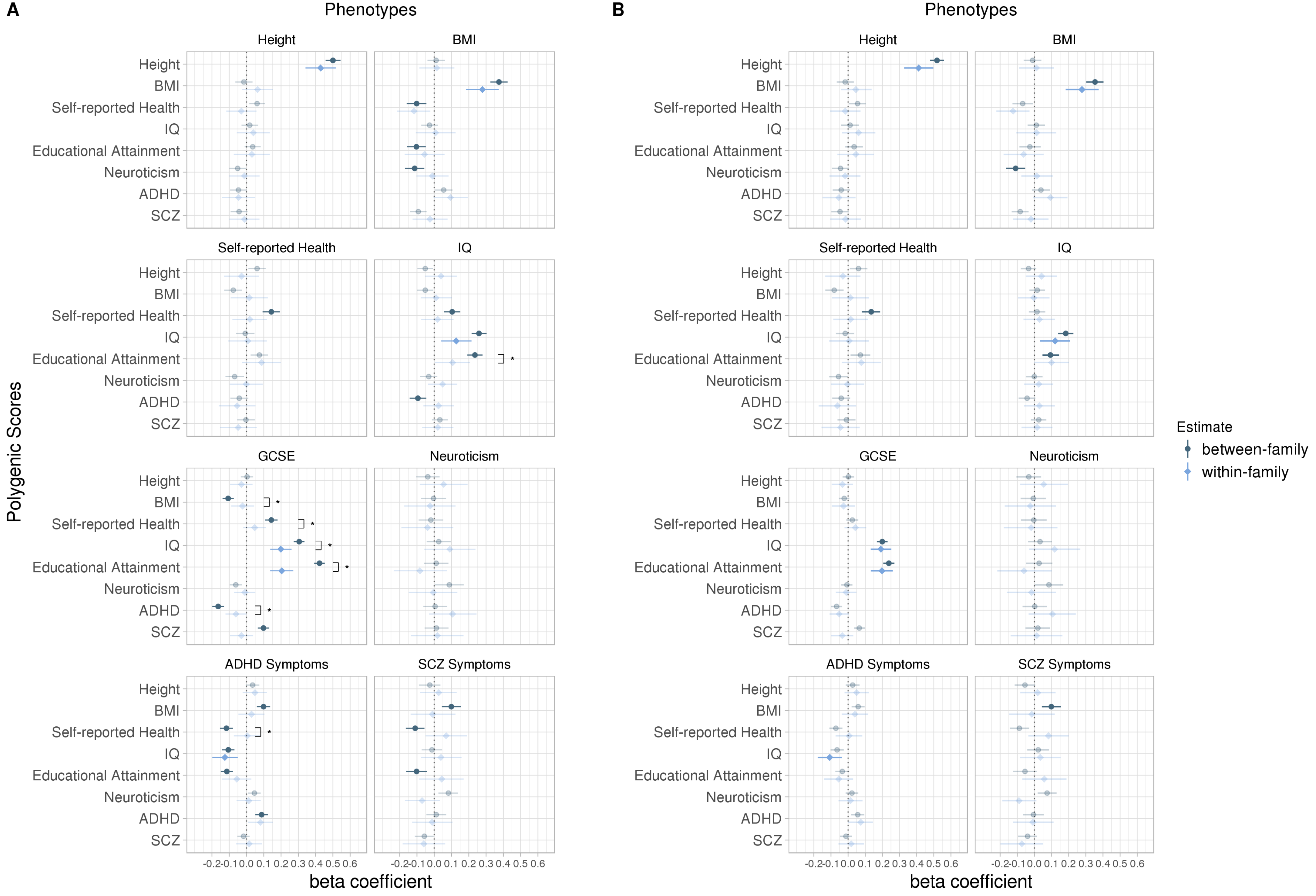

**Note.** Within- and between-family prediction estimates of eight developmental outcomes in opposite-sex twin pairs using eight polygenic scores before (A) and after (B) statistical correction for family socio-economic status, based on opposite-sex twin pairs only. Genome-wide Polygenic Scores are presented on the y-axis, predicting each of the eight phenotypic traits. Error bars are 95% bootstrap percentile intervals based on 10,000 bootstrap samples. Opaque estimates indicate statistical significance at the false discovery rate corrected threshold of $p < 0.01$. Brackets indicate a significant difference between within- and between-family prediction estimate based on permutation testing with 100,000 iterations, and only significant differences are shown where at least one of the estimates is significant at the false discovery rate corrected threshold of $p < 0.01$ (for all prediction estimates and statistical significance, see Supplementary Tables S9 and S10). The dotted line represents a beta coefficient of zero. BMI = Body Mass Index; IQ = Intelligence; GCSE = General Certificate of Secondary Education (educational achievement); ADHD = Attention-Deficit/Hyperactivity Disorder; SCZ = Schizophrenia.

#### References

1. Yengo, L., Sidorenko, J., Kemper, K.E., Zheng, Z., Wood, A.R., Weedon, M.N., Frayling, T.M., Hirschhorn, J., Yang, J., Visscher, P.M., et al. (2018). Meta-analysis of genome-wide association studies for height and body mass index in similar to 700 000 individuals of European ancestry. Hum. Mol. Genet. *27*, 3641–3649.

2. McInnes, G., Tanigawa, Y., DeBoever, C., Lavertu, A., Olivieri, J.E., Aguirre, M., and Rivas, M.A. (2018). Global Biobank Engine: enabling genotype-phenotype browsing for biobank summary statistics. Bioinformatics *9*, 1612.

3. Harris, S.E., Hagenaars, S.P., Davies, G., Hill, W.D., Liewald, D.C.M., Ritchie, S.J., Marioni, R.E., Sudlow, C.L.M., Wardlaw, J.M., McIntosh, A.M., et al. (2017). Molecular genetic contributions to self-rated health. Int J Epidemiol *46*, 994–1009.

4. Savage, J.E., Jansen, P.R., Stringer, S., Watanabe, K., Bryois, J., de Leeuw, C.A., Nagel, M., Awasthi, S., Barr, P.B., Coleman, J.R.I., et al. (2018). Genome-wide association meta-analysis in 269,867 individuals identifies new genetic and functional links to intelligence. Nat. Genet. *50*, 912–919.

5. Allegrini, A., Selzam, S., Rimfeld, K., Stumm, von, S., Pingault, J.-B., and Plomin, R. (2019). Genomic prediction of cognitive traits in childhood and adolescence. Molecular Psychiatry.

6. Lee, J.J., Wedow, R., Okbay, A., Kong, E., Maghzian, O., Zacher, M., Nguyen-Viet, T.A., Bowers, P., Sidorenko, J., Linner, R.K., et al. (2018). Gene discovery and polygenic prediction from a genome-wide association study of educational attainment in 1.1 million individuals. Nat. Genet. *50*, 1112–1121.

7. Luciano, M., Hagenaars, S.P., Davies, G., Hill, W.D., Clarke, T.-K., Shirali, M., Harris, S.E., Marioni, R.E., Liewald, D.C., Fawns-Ritchie, C., et al. (2018). Association analysis in over 329,000 individuals identifies 116 independent variants influencing neuroticism. Nat. Genet. *50*, 6–11.

8. Demontis, D., Walters, R.K., Martin, J., Mattheisen, M., Als, T.D., Agerbo, E., Baldursson, G., Belliveau, R., Bybjerg-Grauholm, J., Bækvad-Hansen, M., et al. (2019). Discovery of the first genome-wide significant risk loci for attention deficit/hyperactivity disorder. Nat. Genet. *51*, 63–75.

9. Pardiñas, A.F., Holmans, P., Pocklington, A.J., Escott-Price, V., Ripke, S., Carrera, N., Legge, S.E., Bishop, S., Cameron, D., Hamshere, M.L., et al. (2018). Common schizophrenia alleles are enriched in mutation-intolerant genes and in regions under strong background selection. Nat. Genet. *50*, 381–389.
